# The dynamin-related protein Dyn2 is essential for both apicoplast and mitochondrial fission in *Plasmodium falciparum*

**DOI:** 10.1101/2024.03.15.585229

**Authors:** Alexander A. Morano, Wei Xu, Neeta Shadija, Jeffrey D. Dvorin, Hangjun Ke

## Abstract

Dynamins, or dynamin-related proteins (DRPs), are large mechano-sensitive GTPases mediating membrane dynamics or organellar fission/fusion events. *Plasmodium falciparum* encodes three dynamin-like proteins whose functions are poorly understood. Here, we demonstrate that PfDyn2 mediates both apicoplast and mitochondrial fission. Using super-resolution and ultrastructure expansion microscopy, we show that PfDyn2 is expressed in the schizont stage and localizes to both the apicoplast and mitochondria. Super-resolution long-term live cell microscopy shows that PfDyn2-deficient parasites cannot complete cytokinesis because the apicoplast and mitochondria do not undergo fission. Further, the basal complex or cytokinetic ring in *Plasmodium* cannot fully contract upon PfDyn2 depletion, a phenotype secondary to physical blockage of undivided organelles in the middle of the ring. Our data suggest that organellar fission defects result in aberrant schizogony, generating unsuccessful merozoites. The unique biology of PfDyn2, mediating both apicoplast and mitochondrial fission, has not been observed in other organisms possessing two endosymbiotic organelles.

**Highlights:**

- PfDyn2 is essential for schizont-stage development.
- PfDyn2 mediates both apicoplast and mitochondrial fission.
- Deficiency of PfDyn2 leads to organellar fission failures and blockage of basal complex contraction.
- Addition of apicoplast-derived metabolite IPP does not rescue the growth defects.

## Introduction

Malaria remains a significant global health burden, with 249 million cases with 608,000 deaths in 2022^1^.

*Plasmodium falciparum* is the causative agent of the deadliest form of malaria, and the clinical disease is caused by the asexual replication of parasites within human red blood cells. *P. falciparum* harbors two distinct bacterially derived organelles, a single mitochondrion and a single apicoplast. The mitochondrion, tracing its origin to an alpha-protobacterium akin to other eukaryotes, is a key target for antimalarial drugs^2–4^. Clinical interventions such as atovaquone^5–7^, introduced in 2000, and a multitude of inhibitors undergoing pre-clinical or clinical trials^8–12^, disrupt the parasite’s mitochondrial electron transport chain or ability to synthesize pyrimidines. The apicoplast is a unique four-membrane-surrounded structure originating from a cyanobacterium through primary and secondary endosymbiosis^13–16^. It has been the target of antibiotics used to treat malaria since the 1950s^17^ and continues to be an important target for developing novel antimalarials^18–22^. These two organelles house numerous essential pathways, such as synthesis of pyrimidines^23^, iron-sulfur clusters^24^, adenosine triphosphate (ATP)^25^, Heme^26^, lipids^27^, isoprenoids^28^, and coenzyme A (CoA)^29^. These metabolites are required for parasites to progress through their life cycle in both human and mosquito hosts. Despite the wealth of knowledge about their biochemical functions and significance for drug development, the mechanisms underlying the division of these organelles remain unknown. In the 48-hour asexual blood stage (ABS), the mitochondrion and apicoplast transform from small globular structures to large tubular networks in dividing parasites^30–32^. At the end of schizogony^33,34^, the specialized cytokinesis of *Plasmodium* and related parasites, the branched mitochondrion or apicoplast is divided into 16-32 copies and subsequently distributed into all progeny, ensuring that each daughter cell (merozoite) acquires a single copy of each organelle^35,36^. The mitochondria and apicoplast cannot be made *de novo*, and organellar expansion and fission processes are required for generating functional merozoites. Additionally, since *P. falciparum* does not encode FtsZ or other proteins present in the bacterial and chloroplast fission machineries^2^, the parasite must rely on unique organellar fission mechanisms.

Dynamins, or dynamin-related proteins (DRPs), mediate membrane scission or fusion events across various cellular processes, including endocytosis, organelle division, and intracellular trafficking^37^. These large GTPases form helical structures around the membranes through self-oligomerization, generating constrictive force to divide their cellular targets^38^. Apicomplexan parasites, including *Plasmodium spp.,* contain three phylogenetically distinct dynamin-related proteins^39–42^. While previous studies indicated PfDyn1 (PF3D7_1145400) and PfDyn2 (PF3D7_1037500) possess GTPase activity^43^, and suggested PfDyn2 localized to the endoplasmic reticulum (ER), Golgi, and apicoplast, their precise localization and function remain elusive^44–45^.

In this study, employing transgenic parasites, super-resolution, ultrastructure expansion microscopy, and live cell time-lapse imaging, we unveil PfDyn2’s role in mediating both mitochondrial and apicoplast fission. PfDyn2 localizes to both organelles in schizont-stage parasites and orchestrates their division. Conditional depletion of PfDyn2 prevents organellar fission with profound defects in merozoite formation. Our findings illuminate a unique aspect of *P. falciparum* biology - in no other organism that possesses two bacterially-derived organelles does a single DRP govern both organelles’ division. Thus, PfDyn2’s role also identifies it as a potentially valuable target for novel antimalarials.

## Results

### PfDyn2 is essential for asexual stage development

The *P. falciparum* genome encodes three dynamin-related proteins, PfDyn1 (PF3D7_1145400), PfDyn2 (PF3D7_1037500), and PfDyn3 (PF3D7_1218500) (**Figure S1a**). Previous phylogenetic analyses suggested that apicomplexan DRPs form distinct evolutionary clades diverging from other dynamins, including ARC5, the dynamin involved in plastid division in plants and red alga^39,40^. Apicomplexan dynamins lack the Pleckstrin-Homology (PH) domain, which enables lipid-binding, and likely require additional adaptor proteins to bind to membranes. We fused three copies of the hemagglutinin (3HA) epitope to PfDyn2’s C-terminus and produced the parasite line NF54attB-PfDyn2-3HA^apt^ (PfDyn2-3HA^apt^) (**Figure S1b**). As described previously^46,47^, the TetR-DOZI-aptamer system regulates gene expression using a small molecule, anhydrotetracycline (ATc). PfDyn2-3HA^apt^’s genotype was confirmed by PCR (**Figure S1c**) and the tagged protein was expressed at the expected molecular weight (**Figure S1d**).

To evaluate PfDyn2’s subcellular localization, we performed immunofluorescence assays (IFAs). PfDyn2 is expressed in schizont-stage but not ring- or trophozoite-stage parasites (**Figure S2**), as previously reported^45^. Using super-resolution microscopy, in early schizogony, when there are fewer than 5 nuclei and the inner membrane complex (IMC) is first visible, PfDyn2 is undetectable (**Figure 1a, b**). In middle and late schizogony PfDyn2 localizes to the apicoplast and the mitochondrion, with increased intensity at the ends of the organelles and nascent ‘branch points’, where separation would occur later (**Figure 1a, b**). To verify PfDyn2’s organellar localization with improved spatial resolution, we performed ultrastructure expansion microscopy (U-ExM)^48,49^. PfDyn2 localizes primarily to the apicoplast in early segmentation (**Figure 1c**). In middle and late segmentation, however, PfDyn2 localizes to both organelles and forms rings around both the apicoplast and mitochondria (**Figure 1c**).

**Figure 1:**
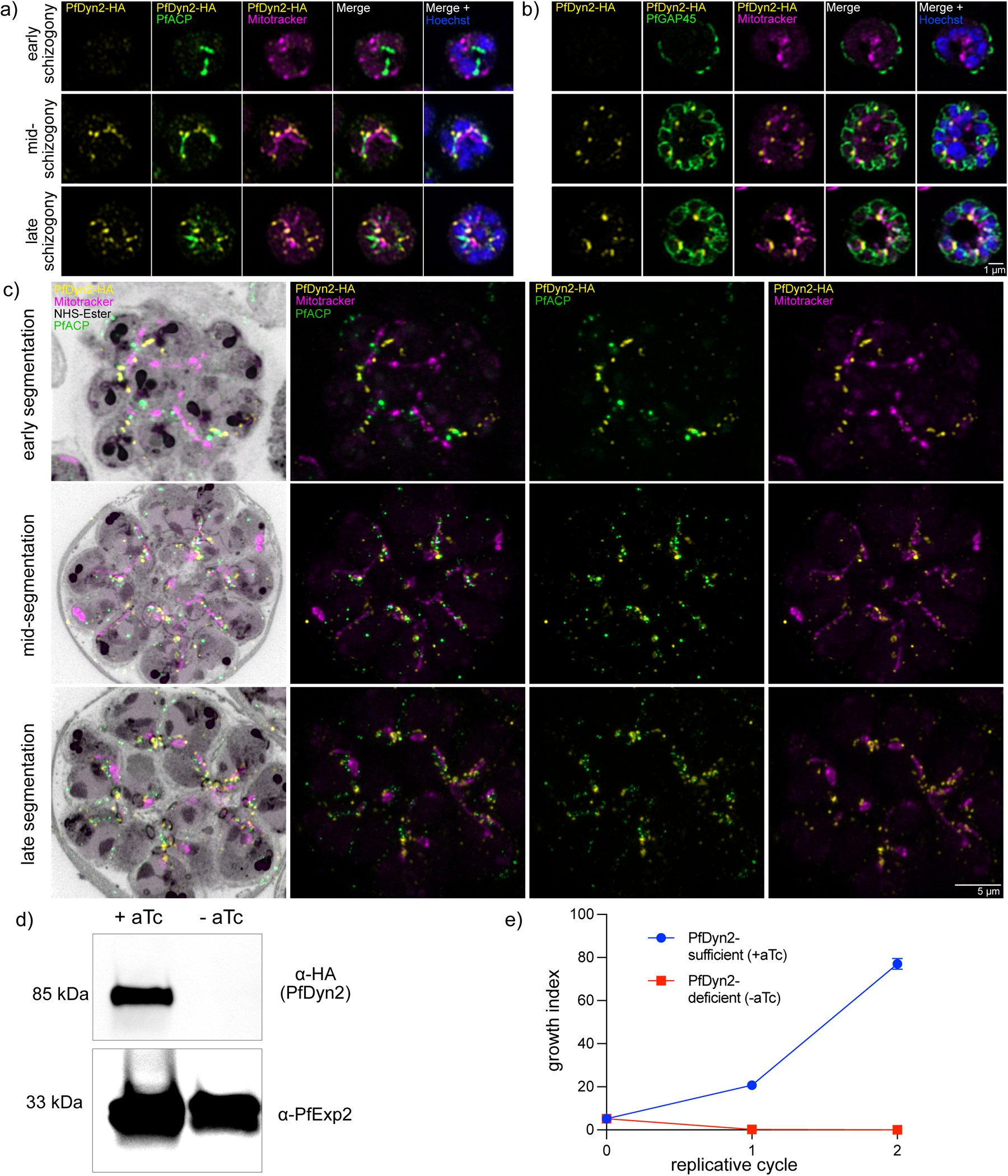
PfDyn2 localizes to the apicoplast and mitochondria and is essential for growth and replication. **a)** Super-resolution microscopy detects PfDyn2’s localization to the apicoplast and mitochondria at different stages of schizogony in PfDyn2-3HA^apt^ parasites. PfDyn2 was probed with anti-HA. The apicoplast was probed with anti-ACP (acyl carrier protein). Mitochondria were labeled with MitoTracker Orange CMTMRos. Nuclei were stained by Hoechst. **b)** Super-resolution microscopy verifies PfDyn2’s localization to mitochondria at different stages of IMC development in the PfDyn2-3HA^apt^ parasites. IMC was probed by anti-PfGAP45. PfDyn2, mitochondria, and nuclei were labelled as shown in (a). **c)** U-ExM of PfDyn2-3HA^apt^ parasites comparing localization of PfDyn2 at different stages of segmentation (early, mid, and late). PfDyn2, the apicoplast, and mitochondria were labeled as shown in (a). Proteins were labeled by AlexaFluor 405 NHS-Ester and shown in grey. **d)** Immunoblot showing extent of PfDyn2 knockdown over 36-h following ATc removal. PfExp2, exported protein 2, was used as a loading control. **e)** Growth curve comparing replicative fitness of PfDyn2-sufficient (+ATc) and -deficient (-ATc) parasites over 96-h (2 replicative cycles). Parasitemia was determined by microscopy examination of Giemsa-stained thin blood smears and was the product of actual parasitemia and splitting factors. Data shown as mean ± SD (n=3). Scale bars for a and b = 1 μm, for c = 5 μm.

To assess PfDyn2’s essentiality, we compared replication rates of PfDyn2-deficient and- sufficient parasites. Following ATc removal, PfDyn2 protein falls below the detection limit after approximately 36 hours (**Figure 1d**), and parasitemia of the PfDyn2-deficient (-ATc) culture fails to increase (**Figure 1e**), establishing that PfDyn2 is essential for asexual growth and replication. To better understand the knockdown phenotype, we performed a detailed time course experiment (**Figure S3**). In a tightly synchronized ring-stage culture (∼ 8-hours post invasion, hpi), we initiated knockdown and examined parasite morphology and PfDyn2 protein levels at specific time points. PfDyn2-deficient parasites first appear morphologically abnormal at the mid-schizont stage (**Figure S3a**). While PfDyn2-sufficient parasites (+ATc) complete segmentation, egress, and reinvade, PfDyn2-deficient parasites (-ATc) arrest as late schizonts and die shortly thereafter. Immunoblot analysis showed that PfDyn2 (+ATc) is not expressed until approximately 40 hpi (**Figure S3b**), corroborating our IFA data (**Figure 1, Figure S2**). These results demonstrate that PfDyn2 is essential for schizont-stage development.

Previous studies suggest that the primary role of the apicoplast during the ABS is to synthesize isoprenoids^28^. Indeed, genetic knockdown of many essential apicoplast genes can be rescued by exogenous isopentenyl pyrophosphate (IPP, 200 µM)^24,50–60^. We thus attempted to rescue PfDyn2-deficient parasites with IPP. We initiated PfDyn2-knockdown for 36 hours then added IPP (200 µM) or ATc and monitored parasite growth for three additional days. We found that exogenous IPP does not rescue PfDyn2 deficiency; IPP treated parasites resembled the -ATc control (**Figure S4a, b**). To ensure our IPP was functional, we performed growth inhibition assays with fosmidomycin^50^, an apicoplast inhibitor that is detoxified by IPP. Parasites (NF54attB, wildtype) were sensitive to fosmidomycin as expected (EC_50_ =1.12 µM), but IPP fully restored their growth under drug treatment (**Figure S4c**). PfDyn2 deficient parasites thus cannot be rescued with isoprenoid supplementation.

### Failure of apicoplast fission upon PfDyn2 depletion

Using the attBxattP integration system^61^, we inserted ACP_L_-mRuby into the genome of PfDyn2-3HA^apt^ parasites. The leader sequence of ACP (ACP_L_), the first 55 amino acids of acyl carrier protein, has been shown to guide fluorescent proteins into the apicoplast matrix^30, 62^. In the PfDyn2-3HA^apt^-ACP_L_-mRuby parasite line, we first verified apicoplast morphologies throughout the ABS (**Figure S5**). In agreement with previous work^30^, we observed that the apicoplast begins elongating and branching to form a network at around 30 hpi. After egress, apicoplasts are distributed into individual merozoites.

Using long-term live cell microscopy, we then tracked PfDyn2-sufficient and -deficient parasites. In PfDyn2-sufficient parasites, we observed sequential apicoplast fission over the course of segmentation (**Figure 2a**, Supp Video 1). The apicoplast first formed a branching structure (**Figure 2a**, top row, 0:40). Next, branches were separated (**Figure 2a**, top row, 1:00-1:40) then individual apicoplasts were separated as they were packaged into merozoites (**Figure 2a**, top row, 2:40-4:20). Upon egress, clear separation between apicoplasts was evident (**Figure 2a**, top row, 4:40). PfDyn2-deficient parasites initially formed branched apicoplasts (**Figure 2a**, bottom row, 0:40-1:20), but few divisions between branches were made (**Figure 2a**, bottom row, 2:40-3:20; Supp Video 2). Upon egress, PfDyn2-deficient apicoplasts collapsed within the residual body (**Figure 2a**, bottom row, 4:40). To quantify this phenotype, we examined >100 parasites per condition which progressed to egress during long-term live cell imaging. PfDyn2-sufficient parasites showed obvious apicoplast segregation immediately pre- or post-egress (95±4.5% per spatial position), whereas no PfDyn2-deficient parasites displayed clear apicoplast segregation (0±0% per spatial position) (**Figure 2b**).

**Figure 2:**
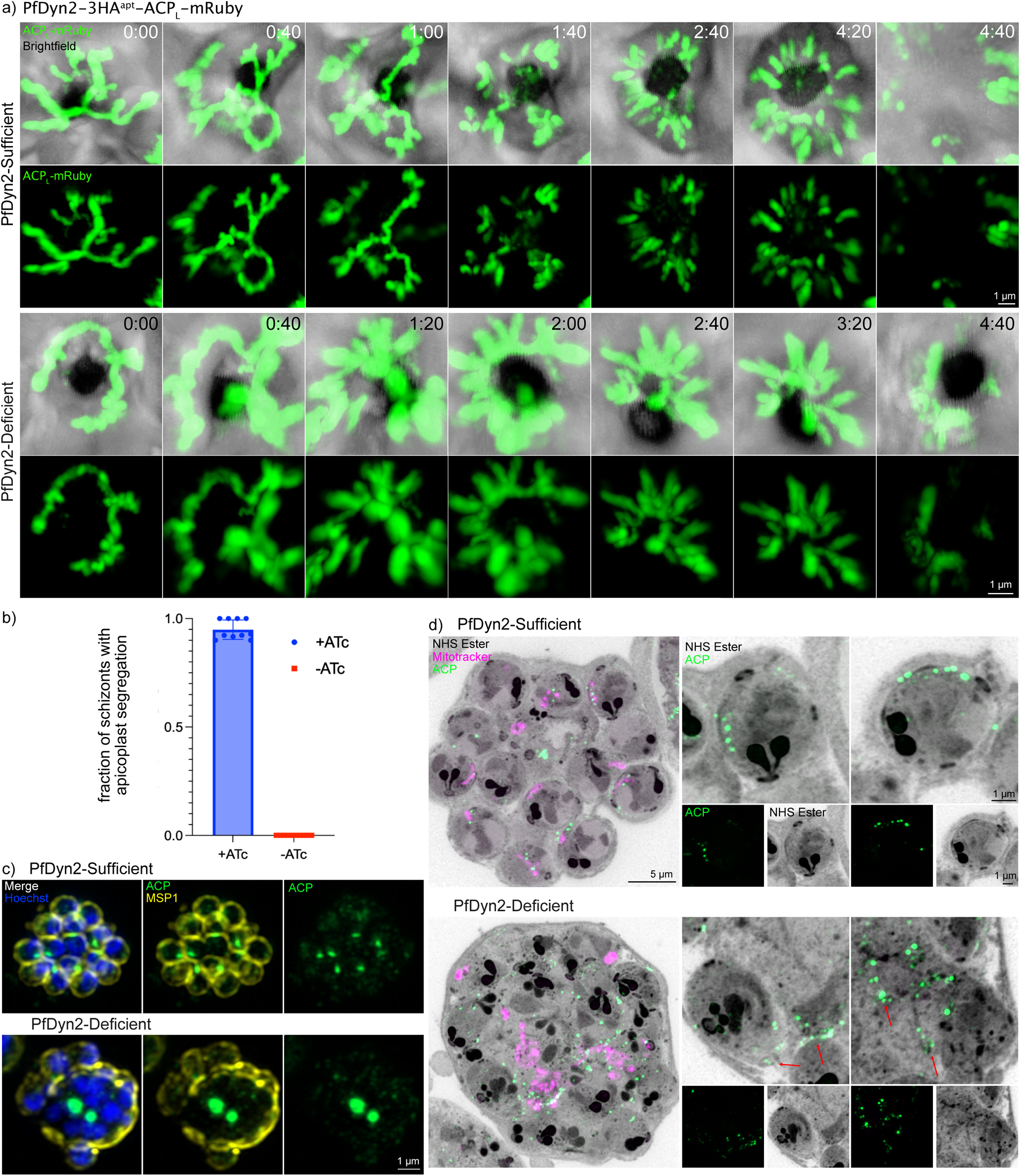
PfDyn2 mediates apicoplast fission. **a)** Images from live cell video microscopy demonstrating apicoplast division in the PfDyn2-3HA^apt^-ACP_L_-mRuby parasites. The top two rows include images from Supp. Video 1 (PfDyn2-sufficient) and bottom two rows include images from Supp. Video 2 (PfDyn2-deficient). Time is displayed as hours:minutes. **b)** Quantification of PfDyn2-3HA^apt^-ACP_L_-mRuby parasites that demonstrated apicoplast segmentation in PfDyn2-sufficient and -deficient conditions under live cell video microscopy. Data shown as mean ± SD with individual values overlaid on bar. **c)** Super-resolution microscopy comparing apicoplast division in PfDyn2-sufficient and -deficient conditions. Late schizont stage parasites were treated with E64 to block egress prior to fixation. PfMSP1(α-PfMSP1) is in yellow, the apicoplast (α-PfACP (acyl carrier protein)) is in green, and nuclei were stained by Hoechst. **d)** U-ExM of PfDyn2-3HA^apt^ parasites comparing apicoplast division in PfDyn2-sufficient and - deficient conditions. Late schizont stage parasites were treated with E64 to block egress prior to fixation. Mitochondria (MitoTracker Orange CMTMRos) are in magenta, apicoplast (α-PfACP) is in green. AF405 NHS Ester was used as a general protein stain and is shown in grey. Maximum projections of Z-stack images are shown on the left. To the right of each max projection are selected merozoites comparing PfDyn2-sufficient apicoplast segmentation and -deficient segmentation. Red arrows point to apicoplasts attached to or leading out from the basal complex. All scale bars = 1 μm except for larger U-ExM images where the scale bar = 5 μm. a-d, Data shown are derived from two biological replicates.

We then examined apicoplast fission in PfDyn2-sufficient and -deficient conditions with greater spatial resolution in fixed cells. We treated parasites with E64 prior to fixation, a cysteine-protease inhibitor that prevents rupture of the RBC plasma membrane^63^, to enrich late schizont-stage parasites that have normally finished organellar fission. Using super-resolution immunofluorescence we labeled the apicoplast with an ACP antibody and the parasite plasma membrane with an antibody against PfMSP1, merozoite surface protein 1^64^. The proper distribution of PfMSP1 to the merozoite surface serves as a marker for normal segmentation. PfDyn2-sufficent schizonts form evenly sized and shaped merozoites, each with a copy of the apicoplast; however, PfDyn2-deficient parasites have a large central mass of apicoplast material surrounded by a disoriented parasite plasma membrane (**Figure 2c**). Using U-ExM and markers for the apicoplast (ACP antibody) and mitochondria (MitoTracker) simultaneously, we saw in PfDyn2-sufficient parasites, the apicoplast successfully separated and aligned with the mitochondria in individual merozoites (**Figure 2d**). In PfDyn2-deficient parasites, the apicoplast showed a lack of division, again with a large central mass, and undivided apicoplast branches connected merozoites through the basal complex (**Figure 2d**). Together, these results strongly indicate that PfDyn2 mediates apicoplast fission.

### Failure of mitochondrial fission upon PfDyn2 depletion

We then inserted Strep II-mNeonGreen-Tom22 into the genome of PfDyn2-3HA^apt^ parasites using the attBxattP system, generating the parasite line PfDyn2-3HA^apt^-StrepII-mNeonGreen-Tom22 (PfDyn2-3HA^apt^-mNG-Tom22 in short). In agreement with published work^30^, the mitochondrion displayed expected developmental progression from a globular structure to a branched tubular network during the ABS (**Figure S6**).

Using long-term live cell microscopy, we tracked PfDyn2-sufficient and -deficient parasites. PfDyn2-sufficient parasites demonstrated expected patterns of mitochondrial branching and sequential cleavage, with a clear separation of individual mitochondria and merozoites upon egress (**Figure 3a**, top rows, Supp video 3). PfDyn2-deficient parasites failed to divide mitochondria. Branching structures were formed, but mitochondria failed to segment beyond one or two branch cuts (**Figure 3a**, bottom rows, Supp video 4). Extended imaging of the depicted schizont after the initial point of egress (**Figure 3a**, bottom rows, 6:00) demonstrated how the failure of mitochondrial segmentation hinders separation of individual merozoites, with the merozoite clump appearing unchanged for over an hour following egress (**Figure 3a**, bottom rows, 7:20). We quantified this phenotype by examining >100 parasites which progressed to egress during imaging. PfDyn2-sufficient parasites (93±3.5% per spatial point) showed obvious mitochondrial segregation immediately pre- or post-egress, whereas very few PfDyn2-deficient parasites did (1.6±2.2% per spatial point) (**Figure 3b**).

**Fig 3:**
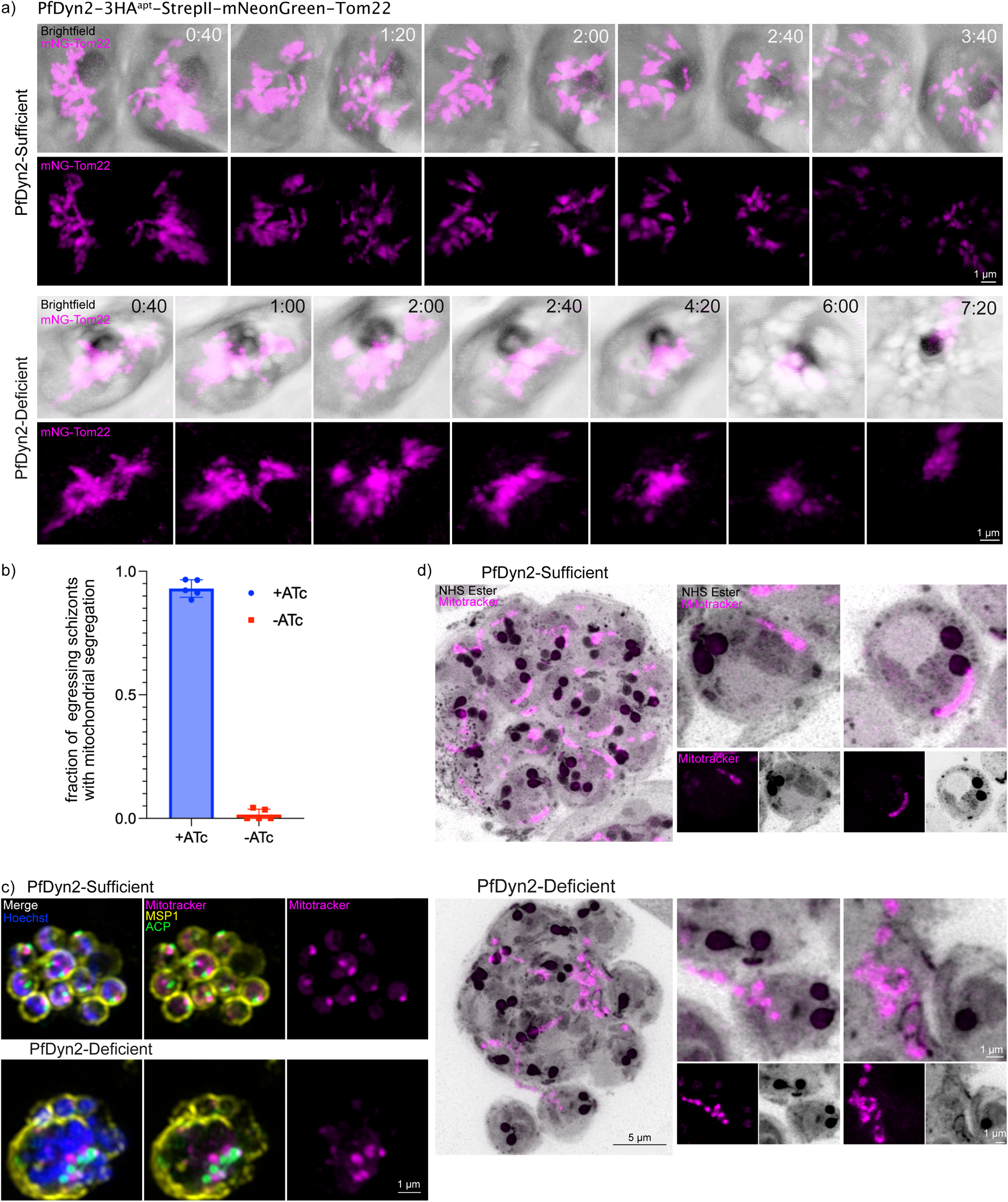
PfDyn2 mediates mitochondrial fission. **a)** Images from live cell video microscopy demonstrating mitochondrial division in the PfDyn2-3HA^apt^-StrepII-mNeonGreen-Tom22 parasites. The top two rows include images from Supp. Video 3 (PfDyn2-sufficient) and bottom two rows include images from Supp. Video 4 (PfDyn2-deficient). Time is displayed as hours:minutes. **b)** Quantification of PfDyn2-3HA^apt^-StrepII-mNeonGreen-Tom22 parasites that demonstrated mitochondrial fission in PfDyn2-sufficient and -deficient conditions under live cell video microscopy. Data shown as mean ± SD with individual values overlaid on bar. **c)** Super-resolution microscopy comparing mitochondrial division in PfDyn2-sufficient and - deficient conditions. Late schizont stage parasites were treated with E64 to block egress prior to fixation. PfMSP1(α-PfMSP1) is in yellow, the apicoplast (α-PfACP) is in green, mitochondria (MitoTracker Orange CMTMRos) are in magenta. Nuclei were stained by Hoechst. **d)** U-ExM of PfDyn2-3HA^apt^ parasites comparing mitochondrial division in PfDyn2-sufficient and -deficient conditions. Late schizont stage parasites were pretreated with E64 to prevent egress. Mitochondria (MitoTracker Orange CMTMRos) are in magenta. Maximum projections of Z-stack images were shown on the left. To the right of each max projection are selected merozoites comparing mitochondrial fission in PfDyn2-sufficient and -deficient conditions. All scale bars = 1 μm except for the larger U-ExM images where the scale bar = 5 μm. a-d, Data shown are derived from two biological replicates.

We next used super-resolution and U-ExM to examine mitochondrial fission at increased resolution in fixed cells. PfDyn2-sufficient parasites’ mitochondria separated such that each merozoite contained a single mitochondrion (**Figure 3c, d**). In PfDyn2-deficient parasites, however, the mitochondrial mass failed to separate, with a central mass of mitochondrial material in the residual body (**Figure 3c, d**). Some merozoites that lack mitochondria are more separated from the mass, but overall, the agglomeration of merozoites is connected by the mitochondrial mass, corroborating live cell imaging data. We occasionally saw undivided mitochondria adjacent to basal complexes that had failed to fully contract, suggesting that mitochondrial division defects can cause segmentation failure (**Figure 3d**). Altogether, these results strongly suggest that PfDyn2 mediates mitochondrial fission.

### Segmentation defects upon PfDyn2 depletion

Schizogony is a highly coordinated, complex process that involves organellar fission as well as the division of one mother cell into 16-32 daughters^33,35^. The inner membrane complex (IMC), a series of flattened vesicles located beneath the parasite plasma membrane, acts as a scaffold, providing structural support to each new merozoite. The basal complex, situated at the posterior end of the newly formed merozoites, guides IMC formation and is thought to provide the contractile force for cell division^65^. Observation of uncontracted basal complexes adjacent to undivided mitochondria led us to examine the IMC and basal complex in PfDyn2-deficient parasites.

We first used U-ExM and an antibody against an IMC-associated alveolin protein, PfIMC1g^66^. PfIMC1g is part of the subpellicular network, which lays directly under the IMC, and it adopts a similar staining pattern to IMC associated proteins like PfGAP45. In PfDyn2-sufficient conditions, PfIMC1g localizes around individual merozoites, each containing one mitochondrion, one nucleus, and a fully contracted basal complex (**Figure 4a**). PfDyn2-deficient parasites had varied types of segmentation defects (**Figure 4a**). The middle row of Figure 4a depicts one phenotype, where many merozoites cannot be distinguished from each other and larger basal complexes are ‘clogged’ with undivided mitochondria. A few amitochondrial merozoites are separated and outlined by PfIMC1g. Thus, there are differences in basal complex size and shape within the same PfDyn2-deficient parasite. The bottom row of Figure 4a shows a slightly different phenotype: each merozoite is outlined by PfIMC1g staining but most mitochondrial material is in the residual body, and basal complexes of merozoites with mitochondria fail to fully contract. Still, in the few amitochondrial merozoites, the basal complex contracts successfully.

**Fig 4:**
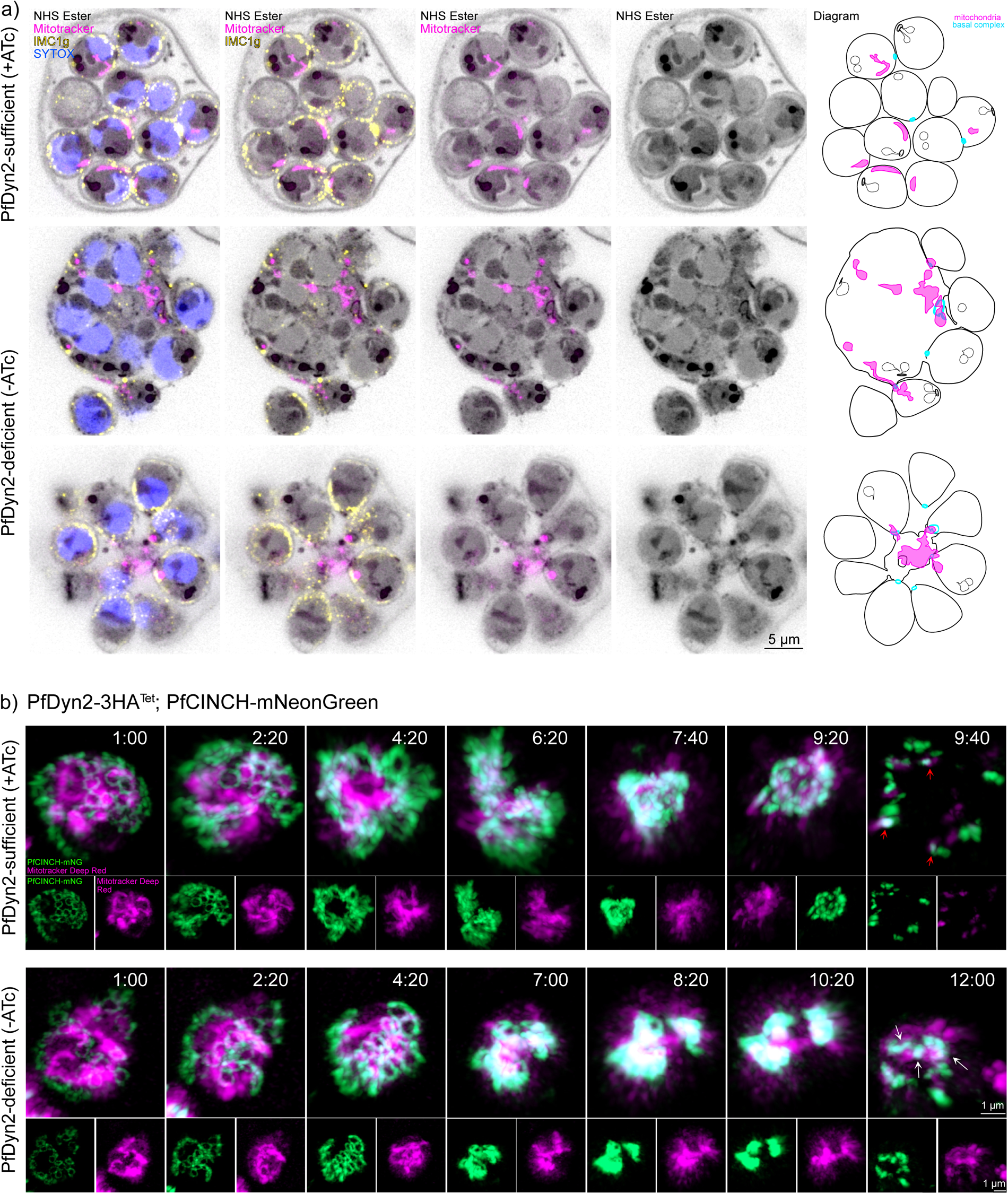
PfDyn2-deficient merozoites fail to complete segmentation. **a)** U-ExM of PfDyn2-3HA^apt^ parasites comparing segmentation in PfDyn2-sufficient and - deficient conditions. Late schizont stage parasites were pretreated with E64 to prevent egress. Maximum projections of Z-stack images (10-20 slices) are shown. The far-right column contains a diagram to help visualize the size and shape of each visible basal complex in the parasite; in the PfDyn2-deficient condition, amitochondrial merozoites are separated from the residual body but merozoites that contain mitochondria are stuck together. The IMC (α-PfIMC1g) is in yellow, the mitochondria (MitoTracker orange CMTMRos) are in in magenta, and nuclei (SYTOX) are blue. AF405 NHS Ester was used as a general protein stain and is shown in grey. **b)** Images from live cell video microscopy in the PfDyn2-3HA^apt^-PfCINCH-mNeonGreen parasites comparing basal complex contraction in PfDyn2 sufficient and -deficient conditions. The top rows include images from Supp. Video 5 (PfDyn2-sufficient) and the bottom rows include images from Supp. Video 6 (PfDyn2-deficient). The basal complex (PfCINCH-mNG) is in green, the mitochondria (MitoTracker Deep Red) are in magenta. Time is displayed as hours: minutes. In PfDyn2-sufficient conditions, upon egress (time point 9:40) the mitochondria separate along with a contracted PfCINCH-basal complex ring (red arrows). In PfDyn2-deficient conditions, upon egress (time point 12:00), PfCINCH rings remain stuck on the mitochondria threaded through them preventing complete contraction (white arrows). Scale bars in A = 5 μm; scale bars in B = 1 μm. a-b, Data shown are derived from two biological replicates.

To observe this process in live parasites, we tagged the basal complex protein PfCINCH^67^ with mNeonGreen in the PfDyn2-3HA^apt^ background and generated the parasite line PfDyn2-3HA^apt^-PfCINCH-mNeonGreen. We then cultured synchronized parasites with and without ATc and performed long-term live cell microscopy after treatment with MitoTracker. In PfDyn2-sufficient parasites, basal complexes synchronously contract around the branched mitochondria as the mitochondria separate (**Figure 4b**, top rows; supp video 5). In PfDyn2-deficient parasites, basal complex initiation and expansion are not affected, but the basal complex fails to fully contract. Mitochondrial staining remains in the center of the schizont; upon egress, mitochondrial branches do not separate, and PfCINCH rings remain threaded onto intact mitochondria (**Figure 4b**, bottom rows; supp video 6).

## Discussion

In this work, we characterize PfDyn2, a novel dynamin-related protein. PfDyn2 localizes to the apicoplast and mitochondria, starting in mid-schizogony and is essential for asexual replication. PfDyn2-deficient parasites could not be rescued with IPP(s). Using super-resolution long-term live cell and ultrastructure expansion microscopy we demonstrate that PfDyn2 is required for apicoplast and mitochondrial fission, as PfDyn2-deficient parasites cannot separate these organelles.

Of the three dynamin-related proteins in *P. falciparum*, PfDyn1 is thought to play a role in hemoglobin uptake^43,68^, PfDyn2 is required for apicoplast and mitochondrial division (this study), and PfDyn3 remains unknown. Immunoprecipitation of basal complex proteins^69^ detected the presence of PfDyn2, indicating PfDyn2 could be involved in basal complex contraction or performing the final scission of merozoites in lieu of ESCRT-III machinery^70–72^, which apicomplexan parasites lack. However, in some PfDyn2-deficient parasites, we observed that amitochondrial merozoites which separate from the schizont have fully contracted basal complexes (**Figure 4a**), suggesting that PfDyn2 is not directly involved in basal complex contraction. PfDyn2 is likely present in basal complex co-immunoprecipitation data because organellar fission occurs as the basal complex contracts and PfDyn2, on the outer membrane of both organelles, interacts with the basal complex close by. Although the three dynamin orthologs are present in *Toxoplasma gondii*, a related apicomplexan parasite, their roles appear different. TgDrpA, the homolog of PfDyn2, is required for apicoplast division, but its depletion does not significantly impact mitochondrial fission^39^, a clear difference between *Plasmodium* and *Toxoplasma*. TgDrpB is involved in secretory organelle biogenesis^40^, and TgDrpC is required for mitochondrial division^42^, with TgDrpC-deficient parasites exhibiting interconnected mitochondria like PfDyn2-deficient schizonts. TgDrpC was also suggested to play broader roles in cytokinesis and organellar biogenesis beyond mitochondrial fission^41^. Our work demonstrates that even orthologous dynamin-related proteins have distinct biological functions in different apicomplexan parasites such as *Plasmodium* and *Toxoplasma*.

Despite having identified the dynamin-related proteins providing the force for organellar fission in both *Plasmodium* and *Toxoplasma*, no homologs to known receptors or recruitment proteins (like Mff or Mid49/51) have been identified in either organism^2^. PfFis1, a *Plasmodium* homolog of a mitochondrial outer membrane adapter protein, is dispensable^73^, as is the *T. gondii* homolog TgFis1^74^, though both localize to the mitochondria. The dispensability of PfFis1/TgFis1 suggests that apicomplexan parasites have additional, divergent mitochondrial outer membrane adaptor proteins. In *Plasmodium*, these adaptor proteins could be common to the apicoplast and mitochondria because PfDyn2 acts on both, making them less likely to be direct homologs of known mitochondrial outer membrane adaptor proteins of model organisms. Further studies, like applying a proximal-biotinylation approach to PfDyn2, could identify putative adaptor proteins common to both endosymbiotically-derived organelles and adaptor proteins unique to each one. In addition to PfDyn2 and receptor proteins yet to be found, a previous study revealed the role of actin in apicoplast fission and cytokinesis^75^. Interestingly, actin depletion only impairs apicoplast fission but not mitochondrial fission, indicating that although PfDyn2 is shared, distinct molecular players are needed to mediate apicoplast fission and mitochondrial fission in *P. falciparum*.

Via U-ExM, PfDyn2 appears to interact with the apicoplast before the mitochondrion (**Figure 1**), which aligns with the organellar fission cascade of late segmentation, where the apicoplast divides approximately 1-2 hours ahead of the mitochondrion^30^. However, the proximity of the two organelles, even in U-ExM, makes conclusive determination difficult. Utilization of higher-resolution microscopy techniques or implementation of Pan-ExM^76^ may demonstrably prove an order of recruitment for PfDyn2. However, we currently cannot rule out the possibility that PfDyn2 binds the apicoplast and mitochondria simultaneously with other factors enabling division of the apicoplast before the mitochondrion.

PfDyn2-deficient parasites demonstrate that preventing organellar fission in *P. falciparum* has severe consequences for segmentation. In PfDyn2-deficient conditions, the basal complex fails to fully contract because undivided organelles are stuck inside it, a conclusion bolstered by the fact that, in some merozoites that lack mitochondria, the basal complex is fully contracted at the end of segmentation (**Figure 4**). IMC distribution is impacted by this as well: failure of mitochondrial and apicoplast division leads to the IMC encircling an undifferentiated mass of cellular material. In these parasites, only a few amitochondrial merozoites are individually packaged. Thus, since the branched mitochondrial and apicoplast structures during schizogony are centered in the residual body, failure of organellar fission prevents segmentation even though basal complex integrity is not compromised. In contrast, basal complex contraction defects do not seem to impact organellar fission – parasites deficient in PfCINCH, required for basal complex contraction, still possessed segmented apicoplast and mitochondria, despite the accumulation of cell materials in large “megazoites”^67^. The IMC and basal complex thus do not actively divide the organelles in *Plasmodium* and their defects do not prevent organellar fission.

The biology of *Toxoplasma* proves an interesting contrast. TgDrpA is required for apicoplast fission, and it was hypothesized that the force generated by daughter bud growth – i.e., by the expansion and contraction of the basal complex – constricted the apicoplast while TgDrpA performed the final separation^39,77^. Parasites defective in TgMORN1, an essential component of *Toxoplasma*’s basal complex^78^, fail to both constrict the basal complex and divide their apicoplast^77^. Interestingly, the TgMORN1 knockout had a milder segregation defect for the mitochondrion (a lack of abscission) compared to the apicoplast, suggesting differential involvement of the basal complex in division of each organelle. In *T. gondii*, the mitochondrion interacts with the pellicle throughout its lifecycle, and the apicoplast makes contact with the pellicle specifically during endodyogeny^79,80^. It makes sense then that basal complex contraction is required for apicoplast division and the final stage of mitochondrial fission in *Toxoplasma*^81^. The fact that *Plasmodium* mitochondria and apicoplast seem to lack contacts with the cytoskeleton during their growth and division may explain why mitochondrial and apicoplast fission defects impact segmentation in *Plasmodium*, but not the other way around^35,82^. Since these endosymbiotically derived organelles don’t form contacts with the cytoskeleton, their division does not rely on cytoskeletal or basal complex integrity but utilizes a different set of proteins^82^. Hence, we observe different apicoplast and mitochondrial phenotypes in PfCINCH-deficient schizonts^67^ compared to TgMORN1-deficient tachyzoites^77^.

In summary, PfDyn2 is essential and localizes to and is required for the division of both the apicoplast and mitochondrion. To the best of our knowledge, for the first time, we utilize novel live cell imaging to visualize apicoplast and mitochondrial division defects and show that segmentation cannot be completed in the absence of functional organellar fission. Though additional studies are needed to identify adaptor proteins and characterize pathways of mitochondrial and apicoplast division in *P. falciparum*, we have already uncovered data that suggests endosymbiotic organellar division in *Plasmodium* occur via a significantly different mechanism than other eukaryotes including the closely related apicomplexan parasite, *Toxoplasma gondii*.

## STAR Methods

Supplemental Experimental Procedures include detailed experimental protocols and oligonucleotide sequences.

1, Plasmid construction. Details of plasmid construction are illustrated in Supplementary Experimental Procedures.

2, Parasite culture, transfection and knockdown studies. These procedures were carried out as previously described^83–85^. Minor modifications are shown in Supplementary Experimental Procedures.

3, Immunoblot analysis. Sample preparation and antibody titers are described in Supplementary Experimental Procedures.

4, Immunofluorescence with an epifluorescence microscope (Nickon Ti), a super-resolution microscope, and Ultrastructure expansion microscopy. Detailed procedures are described in Supplementary Experimental Procedures.

5, Live cell imaging. The long-term live cell imaging experiments were carried out as previously described^69^. Some minor modifications are shown in Supplementary Experimental Procedures.

## Author Contributions

A.M. Investigation, Visualization, Writing – original draft. W.X. Investigation. N.S. Investigation. J.D.D. Formal Analysis, Supervision, Funding Acquisition, Writing – editing and reviewing. H.K. Conceptualization, Investigation, Supervision, Funding acquisition, Writing – original draft. All authors have reviewed and edited the final draft.

## Acknowledgements

We thank Sean Prigge and Scott Lindner for providing the rabbit anti-ACP antisera, Julian Rayner for providing the rabbit anti-PfGAP45 antisera, Anthony Holder for providing the mouse anti-PfMSP1 antibody, and Joshua Beck for providing the NF54attB line. We are grateful for the former members of the Ke laboratory, Dr. Liqin Ling, Rachel Reviello, Dr. Mulaka Maruthi, Dr. Swati Dass, in helping us make plasmid constructs. We thank Ikechukwu Nwankwo for technical assistance. We are grateful for the constructive discussions held with the members of the Center for Parasitology at Drexel University College of Medicine. The study was supported by the National Institutes of Health grants, including R21AI156735 (H.K.), R01 AI169648 (J.D.D.), and F31 AI157041 (A.A.M.).

## Supplemental Experimental Procedures

**Table S1.**
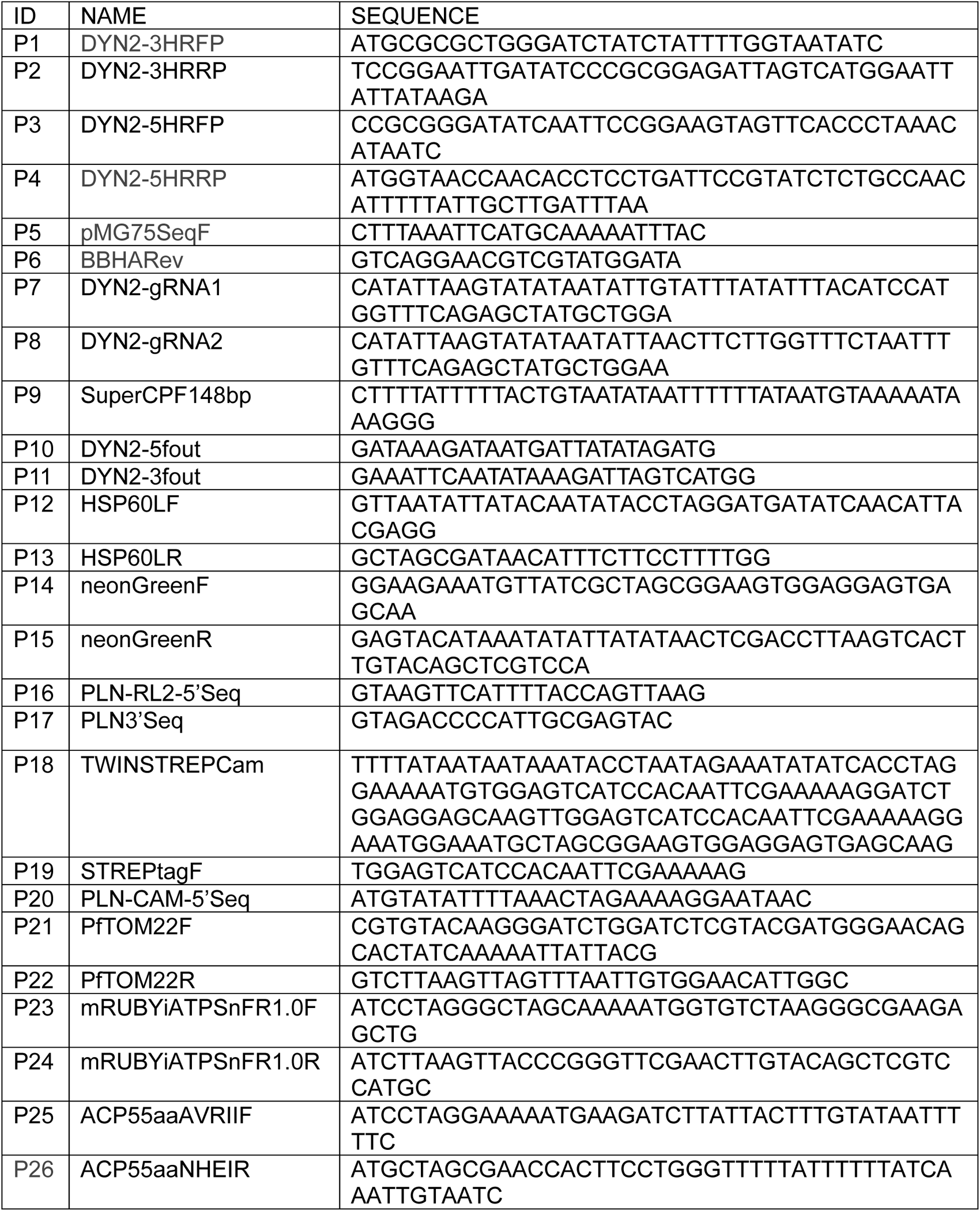
Primers and oligoes used in the study.

### 1, Plasmid construction

#### 1) Conditional knockdown of PfDyn2 using the TetR-DOZI-aptamer system^1,2^

The two homologous regions of PfDyn2 (PF3D7_1037500, 3HR, 5HR) were PCR amplified from genomic DNA using primers P1/P2 (3HR) and P3/P4 (5HR). The DNA fragments were sequentially cloned into the pMG75-BSD-3HA construct^3^ and verified by Sanger sequencing using vector primers (P5 and P6). These procedures produced pMG75-BSD-PfDyn2-3HA, which was linearized with EcoRV prior to transfection. Two gRNA coding sequences (P7, P8) were selected from near the end of the PfDyn2 genetic locus by Eukaryotic Pathogen CRISPR guide RNA/DNA Design Tool (grna.ctegd.uga.edu). They were individually cloned into the NFCas9 plasmid with Infusion as described previously^4^, yielding two gRNA expressing plasmids. The gRNA sequences were verified by Sanger sequencing (P9).

#### 2) Label the mitochondria with mNeonGreen (mNG)

We first tried to express the mNG protein in the mitochondrial matrix with the help of a mitochondrial leader sequence from HSP60^5^. We amplified the leader sequence of HSP60 (the first 68 aa) from genomic DNA using primers P12/P13 and the mNeonGreen fragment from pM2GT-HSP101-mNeonGreen using primers P14/P15. The two PCR amplicons were cleaned and fused together with the pLN-RL2-BSD-RL13-3HA vector that was digested with AvrII and AflII, resulting in the plasmid, pLN-RL2-BSD-HSP60L-mNG. The HSP60L-mNG sequence was verified by Sanger sequencing using vector primers (P16, P17). The insert, HSP60L-mNG, was further cloned into pLN-Cam-BSD-yDHODH-GFP^6^ by restriction sites AvrII and AflII, resulting in another plasmid, pLN-Cam-BSD-HSP60L-mNG. Transgenic parasites were made with pLN-RL2-BSD-HSP60L-mNG and pLN-Cam-BSD-HSP60L-mNG plasmids, but their fluorescence was either too dim (RL2) or too strong (Cam), not suitable for labeling the mitochondrion (data not shown).

We then tried to label the mitochondria with mNG by fusing it to the N-terminus of Tom22. Tom22 is a tailed anchor protein and localizes to the mitochondrial outer membrane^7^. We also added a twin Strep tag upstream of mNG-Tom22, which can facilitate mitochondrial purification in future studies. Starting from pLN-Cam-BSD-HSP60L-mNG, we used Infusion to replace the HSP60L sequence with a synthetic gene block containing the Strep II sequence (P18), yielding pLN-Cam-BSD-StrepII-mNG. Colony PCR was performed to select positive clones (P19/P17). The correct sequence of StrepII-mNG was verified by sequencing (P20). Next, pLN-Cam-BSD-StrepII-mNG was further digested with BsrGI and AflII and ligated with Tom22, which was amplified from genomic DNA using primers P21/P22. The correct Tom22 sequence was verified by Sanger sequencing (P17). These procedures yielded pLN-Cam-BSD-StrepII-mNG-Tom22. To switch the selectable marker from BSD to hDHFR, StrepII-mNG-Tom22 was excised by AvrII and AflII and subsequently cloned into two pLN plasmids, pLN-RL2-hDHFR-RL13-3Myc^8^ and pLN-Cam-hDHFR-VP1-3Myc^3^. These procedures produced two plasmids, pLN-RL2-hDHFR-StrepII-mNG-Tom22 and pLN-Cam-hDHFR-StrepII-mNG-Tom22. Transfections of both plasmids were conducted; however, only parasites transfected with pLN-RL2-hDHFR-StrepII-mNG-Tom22 had viable parasites and proper labeling of mitochondria.

#### 3) Label the apicoplast with mRuby

We PCR amplified mRuby from PM2GT-mRuby-Hsp101 using primers P23/P24 and subsequently cloned it into pLN-RL2-hDHFR-RL13-3Myc^8^ using AvrII and AflII, resulting in pLN-RL2-hDHFR-mRuby. The leader sequence of ACP (first 55 aa) was PCR amplified from genomic DNA using primers P25/P26 and cloned into pLN-RL2-hDHFR-mRuby by AvrII and NheI, yielding pLN-RL2-hDHFR-ACP_L_-mRuby. The correct sequence of ACP_L_-mRuby was verified by Sanger sequencing (P16).

### >2, Parasite culture and transfection

Wild type *P. falciparum* strains, D10 and NF54attB, were used in the study. Parasites were grown in complete RPMI-1640 media supplemented with Albumax I (0.5%) and human O^+^ RBCs, as previously described^8^. Transfections were performed in ring stage parasites (> 5% parasitemia) with either linearized or circular plasmids (∼ 50 µg) using a BioRad electroporator. Post electroporation, the cultures were kept in a low oxygen atmosphere (90% N2, 5% CO2, 5% O2) and added with proper drug selections, e.g., blasticidin (2.5 µg/mL, InvivoGen), WR99210 (5 nM, Jacobs Pharmaceutical), G418 (125 µg/mL, VWR), or anhydrotetracycline (aTc) (250 nM, Fisher Scientific).

To construct the line PfDyn2-3HA^apt^-PfCINCH-mNeonGreen, 25 μg of PfCINCH-HDR plasmid containing mNeonGreen (pRR208) was linearized by digestion with STUI, purified, and co-transfected with 20 μg of a guide plasmid (pRR99), a construct containing an SpCas9 expression cassette and the PfCINCH-targeting guide RNA. Transfection was performed in the NF54attB-Dyn2-3HA^apt^ line. Parasites were maintained with 2.5 μg/mL Blasticidin and 500 nM Anhydrotetracycline (ATc) from the onset of transfection. One day post transfection, drug pressure for the second marker was applied with 5 nM WR99210.

### 3, Knockdown studies/Western blot analysis

To initiate knockdown studies, the synchronized NF54attB-PfDyn2-3HA^apt^ parasites at the ring stage were thoroughly washed with 1xPBS to remove aTc and diluted in fresh RBCs to receive ATc (+) or (-) media. For regular knockdown studies, thin blood smears and parasite proteins were harvested daily or every two days. For time-course knockdown studies, samples were collected every 4 hours for 6 times, starting at 24-hour post ATc removal. To extract proteins, the parasite cultures were treated with 0.05% Saponin/PBS supplemented with 1x protease inhibitor cocktail (Apexbio Technology LLC), and the pellets were solubilized by 2%SDS/Tris-HCl (65 mM, pH 6.8). The other Western blot procedures followed standard protocols. The blots were incubated with primary antibodies, including the HA probe (mouse, sc-7392, Santa Cruz Biotechnology; 1:10, 000) and anti PfExp2 (rabbit, a kind gift from Dr. James Burns, Drexel University; 1:10, 000). Secondary HRP-labelled antibodies were purchased from ThermoFisher Scientific, including goat anti-mouse (A16078, ThermoFisher Scientific, 1:10, 000) and goat anti-rabbit antibody (31460, ThermoFisher Scientific, 1:10,000). The blots were incubated with Pierce ECL substrates and developed by the ChemiDoc Imaging Systems (Bio-Rad).

### 4, Immunofluorescence analysis (IFA) with Nickon Ti

NF54attB-PfDyn2-3HA^apt^ parasites were tightly synchronized with several rounds of alanine/HEPES (0.5M/10 mM). In various stages (ring, trophozoite, schizont), aliquots of parasite cultures were removed (∼ 50 µL per aliquot). Prior to being fixed with 4% formaldehyde/0.0075% glutaraldehyde, they were either labeled with 60 nM of MitoTracker Red CMXRos (M7512, ThermoFisher) for 30 min or left untreated. After fixation, the cells were permeabilized with 0.25% Triton X-100/PBS, reduced with NaBH4 (0.1 mg/mL), blocked with 3%BSA/PBS. The cells were incubated with primary antibodies, such as the HA probe (mouse, sc-7392, Santa Cruz Biotechnology; 1:300) and/or anti-ACP (rabbit, a kind gift from Dr. Sean Prigge, 1:500). Fluorescently labelled secondary antibodies were purchased from Life Technologies (ThermoFisher Scientific) (anti-mouse or anti-rabbit, 1:300). The samples were stained with DAPI for 10 min (1.5 µg/mL in PBS) and mounted on glass slides in antifade buffer (S2828, ThermoFisher Scientific). Images were captured using a Nickon Ti microscope and processed using the Nickon NIS elements software.

### 5, Immunofluorescence analysis (IFA) with a super-resolution microscope

Schizonts were percoll-purified, resuspended in 100 μL media (with or without 300 nM MitoTracker Orange CMTMRos (M7510, ThermoFisher)) and placed on poly-D lysine treated 10 mm diameter #1.5 coverslips to settle for 25 minutes at 37°C in one well of a 24 well plate. Excess / unbound cells were removed and 300 μL of 4% paraformaldehyde (PFA) in PBS was added for 20 minutes of fixation. PFA was removed and the coverslip was washed 3 times with 1x PBS. 300 μL of 0.1% Triton-X 100 in PBS was then added to the well for 10 minutes to permeabilize the cells. Coverslip was washed 3 times for 3 minutes with 1x PBS following permeabilization. 3% BSA in PBS was added for blocking for one hour, after which relevant primary antibodies resuspended in 3% BSA in PBS were added and the coverslips were incubated overnight at 4°C. Coverslips were then washed 3 times with 1x PBS for 3 minutes again, and secondary antibodies were added for 45 minutes in the dark at room temperature, diluted 1:1000 in 0.5% BSA in PBS. Coverslips were again washed 3 times for 3 minutes with 1x PBS. For 10 minutes, a 1:5000 dilution of Hoechst 3342 in 1x PBS was added to the coverslips. Coverslips were then rinsed with 1x PBS one more time before they were adhered to slides with 5 μL of VectaShield Vibrance (hardening, non-DAPI). IFAs were visualized at least 3-4 hours after mounting. Cells were visualized on a Zeiss LSM 900 with AiryScan 2 for super-resolution microscopy, with a 63X magnification objective with a numerical aperture of 1.4.

Rat anti-HA (used at 1:250) was purchased from Sigma (catalog 11867423001), Rabbit anti-PfGAP45 (used at 1:5000) was a generous gift from Julian Rayner at the Cambridge Institute for Medical Research, Rabbit anti-PfACP (used at 1:1000) was a generous gift from Scott Lindner at Pennsylvania State University. Mouse anti-MSP1 (used at 1:500) was a generous gift from Anthony Holder at the Francis Crick Institute. All secondary antibodies (anti-rat AF488, anti-rat AF647, anti-rabbit AF488) were purchased from Life Technologies and used at 1:1000.

### 6, Ultrastructure Expansion Microscopy^9^

Synchronized Dyn2-3HA^apt^ ATc (±) schizonts were percoll-purified as for standard immunofluorescence. Isolated schizonts were placed on poly-D-lysine coated coverslips for 20-30 minutes at 37°C to settle in one well of a 24 well plate (with 300 nM MitoTracker Orange CMTMRos). Parasites were then fixed with 4% PFA for 20 minutes at 37°C and washed 3X with PBS. Following fixation, the coverslips were incubated with FA/AA (formaldehyde/acrylamide) overnight at 37°C. The next morning, to polymerize the gel, TEMED and APS were swiftly added to a previously made monomer solution containing acrylate, acrylamide, and BIS. The monomer solution was placed as a droplet inside a humid chamber stored at -20°C for 15 minutes preceding gelation. Coverslips were then placed over this drop of the TEMED/APS/Monomer Solution. After 5 minutes of incubation on ice, the chamber was incubated at 37°C for an hour. Coverslips/gels were then placed in 2 mL of denaturation buffer (in one well of a 6 well plate) and incubated with denaturation buffer for 15 minutes with agitation at room temperature. After gel detachment, gels were incubated for 90 more minutes at 95°C in a 1.5 mL Eppendorf tube filled with denaturation buffer. Denaturation buffer was then removed, and gels were placed in a 10 cm dish filled with ddH2O. ddH2O was replaced after 30 minutes, twice. Gels were then incubated in ddH2O overnight. The next morning, gels were washed in 1xPBS 2 times for 15 minutes each then incubated in 3% BSA-PBS for 30 minutes at RT. Blocking buffer was removed and gels were incubated in 1 mL of 3% BSA-PBS with primary antibodies overnight at 4°C. The next day, gels were washed 3 times with 2 mL 0.5% Tween20 in PBS for 10 minutes at room temperature with agitation. Gels were then incubated in 1 mL of PBS with secondary antibodies including NHS-Ester (405, 1:100) protected from light, with agitation at room temperature. After 2.5 hours of secondary antibody incubation, gels were washed 3 more times with 2 mL 1% PBS + 0.5% Tween20 as before, then placed in a 10 cm dish filled with ddH2O. Water was replaced after 30 minutes, twice, and gels were allowed to expand overnight before imaging on a Zeiss laser-scanning confocal 900 with AiryScan 2 for super-resolution microscopy, with a 63X magnification objective and a numerical aperture of 1.4.

### 7, Live Cell Imaging

Parasites were synchronized by means of sorbitol treatment and percoll density centrifugation one or two cycles (2 or 4 days) prior to the experiment. On the day of the experiment, schizonts were percoll-purified and placed in a 5 mL (mm) culture dish with 5 mL of complete media to recover for 30 min. They were then gently spun down and resuspended in 100 µL of media and placed in one quadrant of a concanavalin A-coated iBidi/cellview glass bottom dish. After 30 minutes incubation at 37°C, excess cells were washed off with 3x washes of 1x PBS and each quadrant in use was filled with phenol red-free RPMI with 0.5 mM Trolox added. Parasites were imaged in a 37°C incubation chamber supplemented with 5% CO2; the glass bottom dish was also inset into a heated stage (heated to 37°C) on a Zeiss LSM900 with AiryScan 2. The SR-Multiplex Mode (4Y) was used and images were taken every 20 minutes over 13 hours. For quantification, resulting files were visualized in FIJI. To produce supplementary videos and high-resolution images, resulting files were processed in ARIVIS 4D.

For imaging the PfDyn2-3HA^apt^-PfCINCH-mNeonGreen line, MitoTracker Deep Red FM was added to the media to a final concentration of 10 nM during the settling stage to label mitochondria.

Laser use was dependent on the fluorophore or dye utilized for each experiment. PfDyn2-3HA^apt^-PfCINCH-mNeonGreen imaging (Figure 4), the 488 laser was used at 0.9% laser power and 850 gain and the 647 laser was used at 1.0% laser power with 750 gain. For PfDyn2-3HA^apt^-StrepII-mNeonGreen-Tom22 imaging (Figure 3), the 488 laser was used at 0.9% laser power and 850 gain and the 647 laser was used for transmitted light / brightfield imaging through ESID. For PfDyn2-3HA^apt^-ACP_L_-mRuby imaging (Figure 2), the 561 laser was used at 0.9% laser power and 800 gain and the 647 laser was used for transmitted light / brightfield imaging through ESID.

### 8, Quantification

For Figure 2 (quantification of apicoplast segmentation), 10 fields per condition were taken with 9-15 parasites per field. Parasites were considered “positive” for successful apicoplast division if, at the time point immediately pre-egress or, that time point was unclear, the time point of immediately after egress, there were visible and obviously separated apicoplasts (see Figure 2a for an example of this). In most cases, the resolution was not high enough to determine how many new apicoplasts had been generated, but a binary determination of segmented apicoplasts / nonsegmented apicoplasts could generally be determined. If it was unclear whether a parasite had segmented apicoplasts or not, it was not included in the analysis. Only parasites that egressed during the imaging time course were considered part of the dataset.

For Figure 3 (quantification of mitochondria segmentation), 5 fields per condition were taken with 15-30 parasites per field. Parasites were considered “positive” for successful mitochondrial division if, at the time point immediately pre-egress or, that time point was unclear, the time point of immediately after egress, there were visible and obviously separated mitochondria (see Figure 3a for an example of this). In most cases, the resolution was not high enough to determine how many new mitochondria had been generated, but a binary determination of segmented mitochondria / nonsegmented mitochondria could generally be determined. If it was unclear whether a parasite had segmented mitochondria or not, it was marked as negative (see the comment above). Only parasites that egressed during the imaging time course were considered part of the dataset. Parasites that failed to egress (whether because they were not mature for schizogony or because they died during the imaging process) were excluded from analysis.

## Supplementary Figure Legends

**Figure S1.**
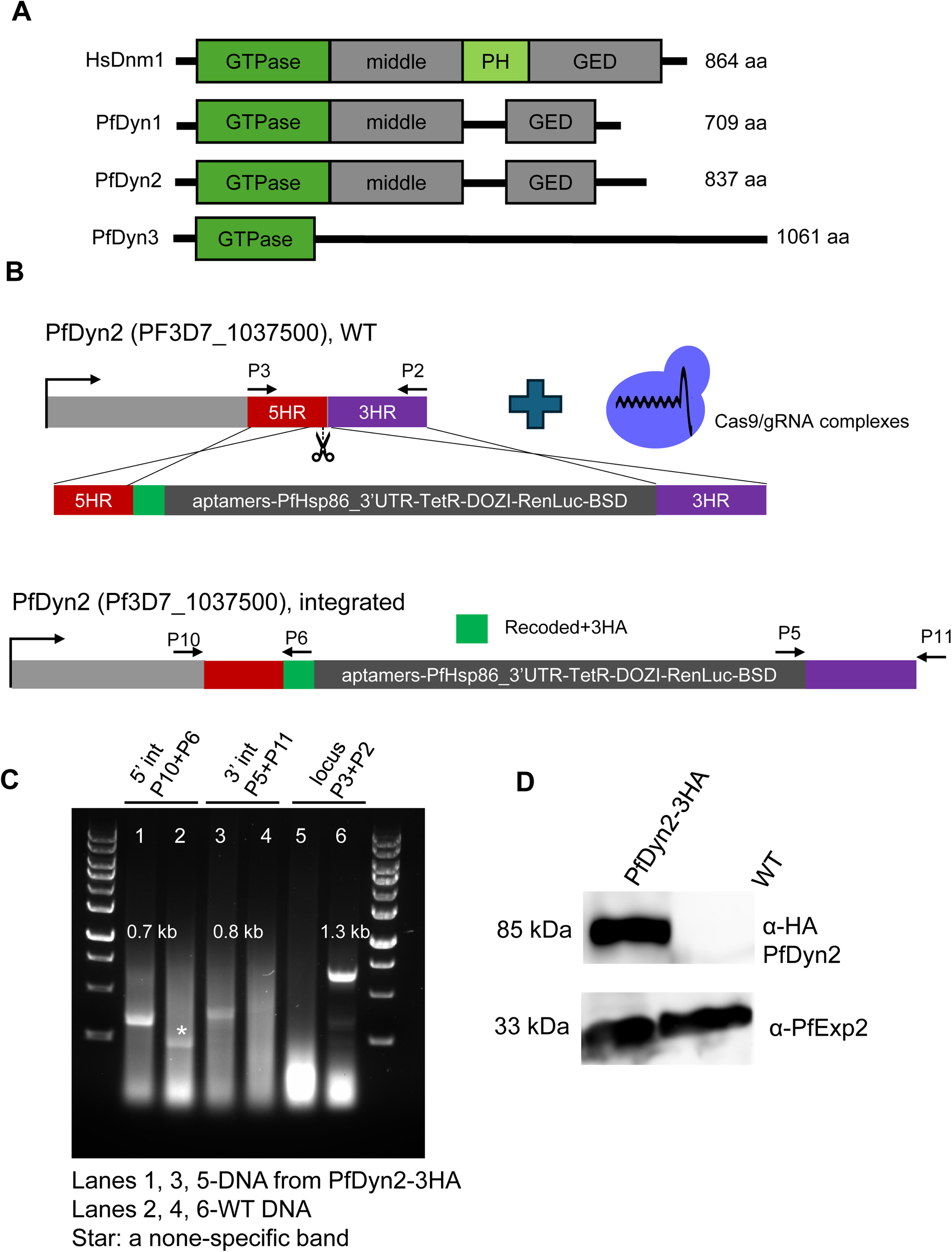
Endogenous tagging of PfDyn2 via CRISPR/Cas9. A, Major domains of the three dynamin like proteins in *P. falciparum* (PfDyn1-3). PfDyn1-3 are divergent from human dynamin 1 and they lack the Pleckstrin homology domain (PH). B, Schematic of CRISPR/Cas9 mediated double crossover recombination strategy. The genetic locus of PfDyn2 (PF3D7_1037500) was added with DNA sequences coding a recoded gRNA sequence, a 3HA tag and elements of the TetR-DOZI-aptamer knockdown system. Primers used for diagnostic PCR in C were indicated. C, Genotyping PCR showing the correct modification of PfDyn2. PCR amplicon sizes were shown. D, Western blot showing the expression of PfDyn2-3HA with a correct molecular weight. WT (wild type) lysate was derived from the parental parasite line, NF54attB. PfExp2 was used as a loading control.

**Figure S2.**
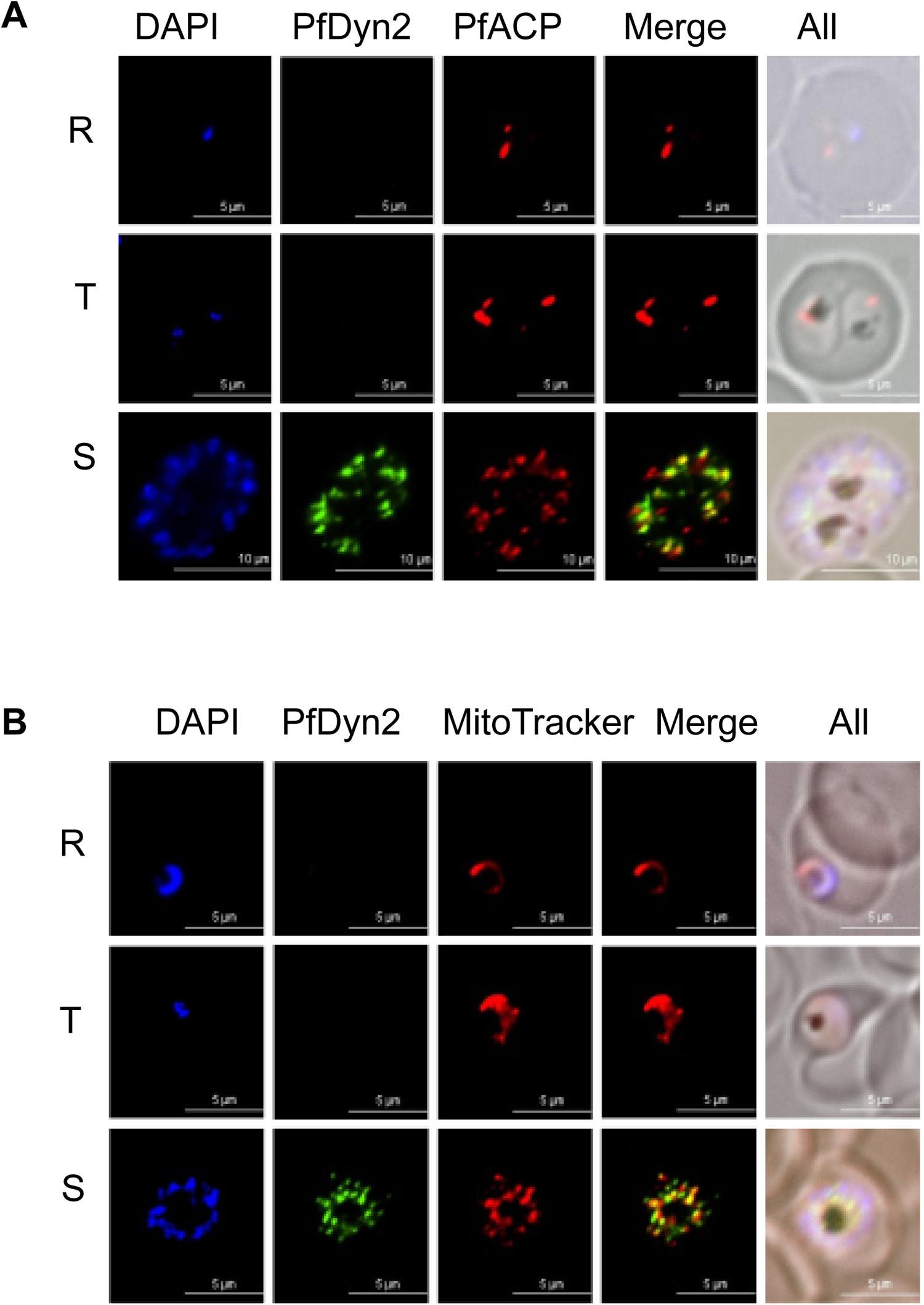
Localization of PfDyn2 across the asexual blood stage. A, Immunofluorescence assays (IFA) showing localization of PfDyn2 to the apicoplast in the schizont stage, but not in the ring or trophozoite stages. DAPI stains nucleus. PfDyn2 was detected by α-HA and FITC-labeled secondary antibody. The apicoplast was labeled with α-ACP, acyl carrier protein, and TRITC-labeled secondary antibody. B, Immunofluorescence assays (IFA) showing localization of PfDyn2 to the mitochondrion in the schizont stage, but not in the ring or trophozoite stages. DAPI stains nucleus. PfDyn2 was detected by α-HA and FITC-labeled secondary antibody. The mitochondrion was labeled with MitoTracker Red CMXRos (60 nM). A-B, images were taken from a Nickon Ti microscope and processed by Nikon NIS elements software.

**Figure S3.**
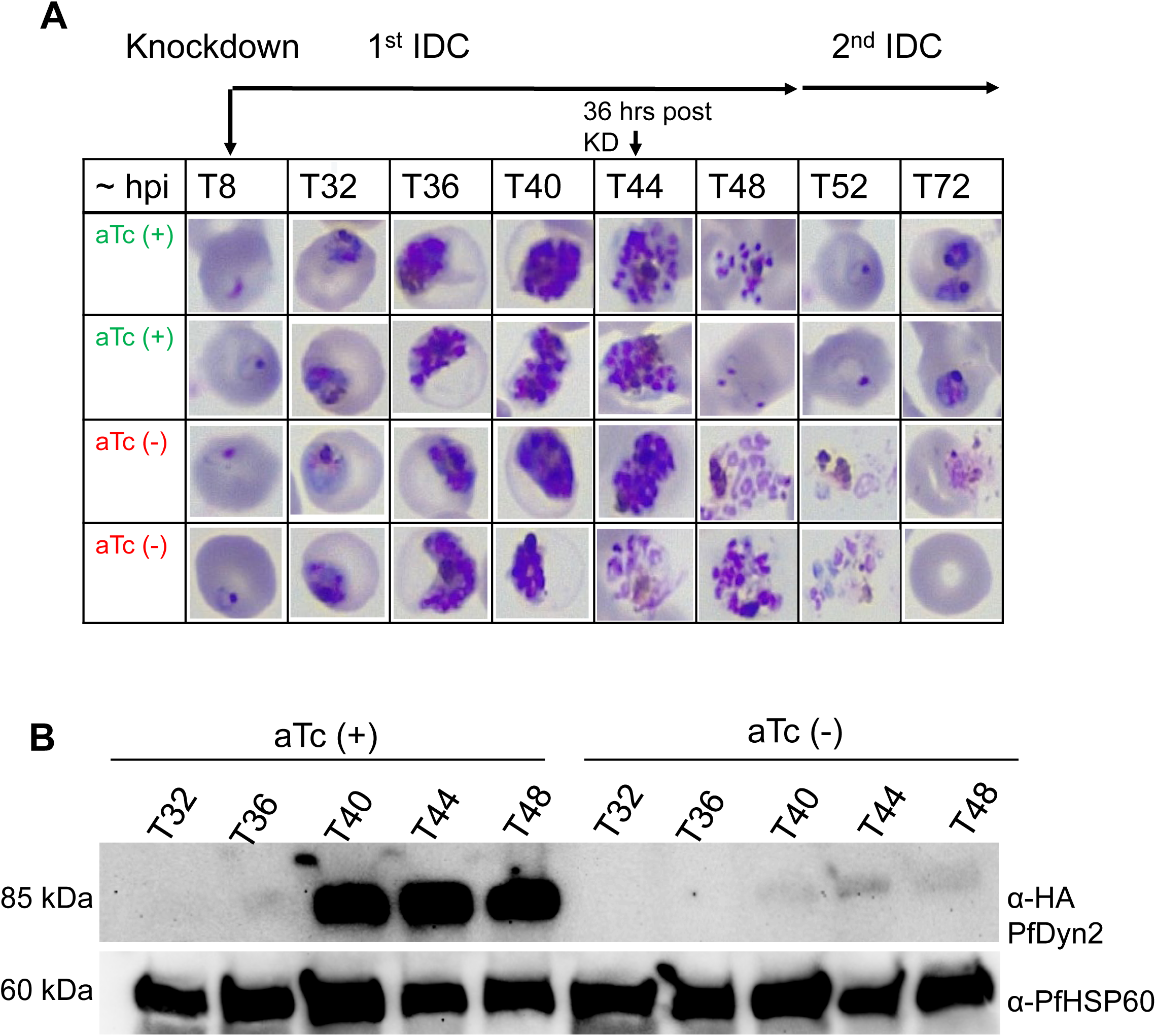
PfDyn2 deficient parasites in a time course experiment. A, Giemsa-stained images showing the morphologies of PfDyn2 sufficient and deficient parasites. ATc was removed from a tightly synchronized ring stage culture. At 36-h post ATc removal, abnormal morphologies in PfDyn2 deficient parasites were visible. B, Western blots showing PfDyn2 expression in PfDyn2 sufficient and deficient parasites. In the presence of ATc, PfDyn2 was detected in the schizont stage. Upon ATc removal, PfDyn2 levels fell below the detection limit. PfHSP60 was used as a loading control (α-PfHSP60, NBP2-12734, Novus).

**Figure S4.**
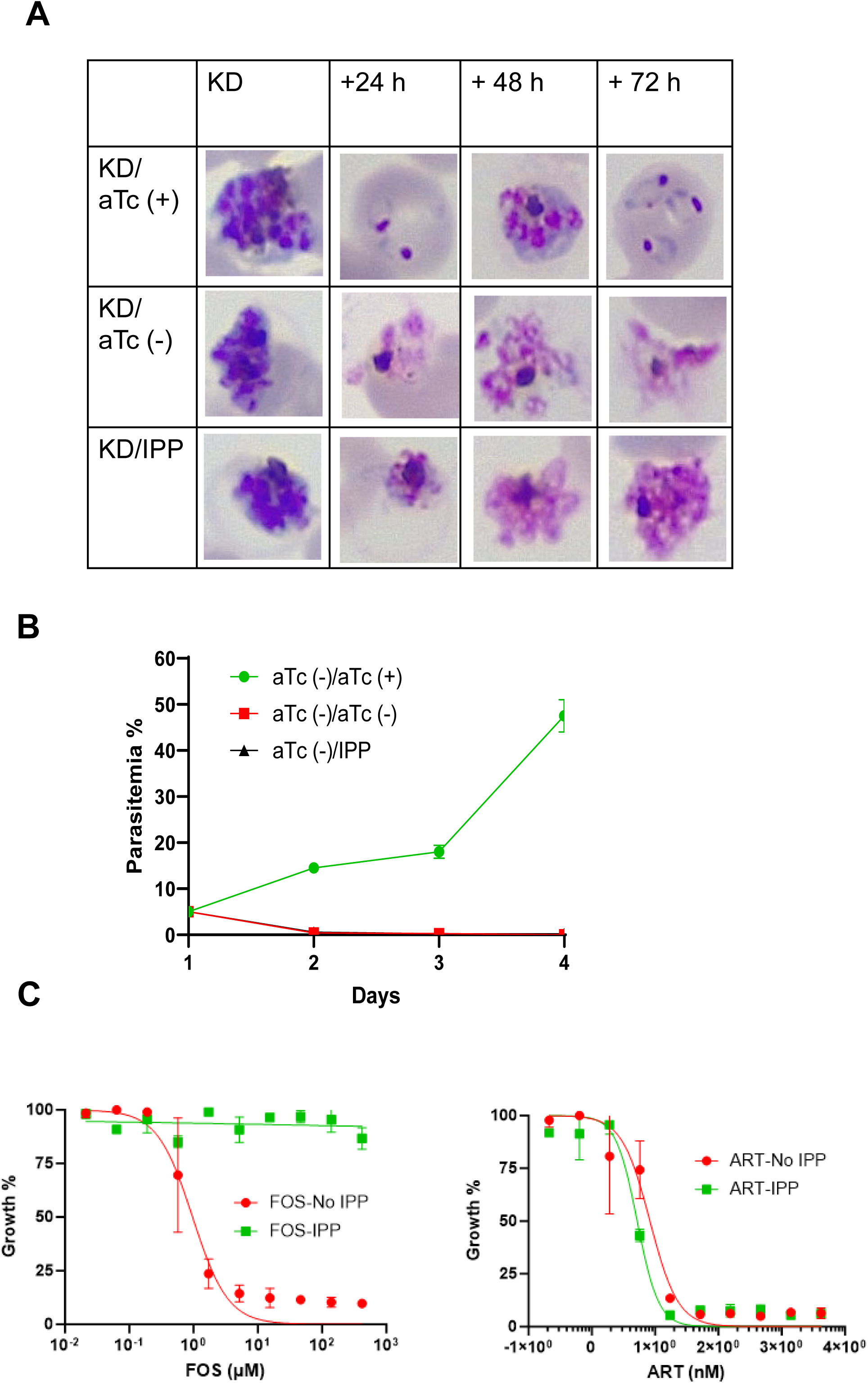
PfDyn2 deficient parasites cannot be rescued with IPP (isopentenyl pyrophosphate, 200 µM). A, Morphologies of PfDyn2 deficient parasites rescued with media supplemented with ATc (+), ATc (-), or IPP (200 µM, IPP001, Isoprenoids, LC). KD, knockdown for 36-h from a tightly synchronized ring stage culture. B, Growth curves of PfDyn2 deficient parasites rescued with ATc (+), ATc (-), or IPP (200 µM). Parasitemia was determined from microscopic counting of at least 1,000 red blood cells in each thin blood smear and is the product of actual parasitemia and splitting factors. Error bars indicate mean ± s.d. of three counts. C, Growth inhibition assays showing toxicity of fosmidomycin, but not artemisinin, can be reversed with IPP (200 µM). SYBR green assays were performed according to the standard 72-h protocol starting from ring stage parasites. Error bars indicate mean ± s.d. of three replicates.

**Figure S5.**
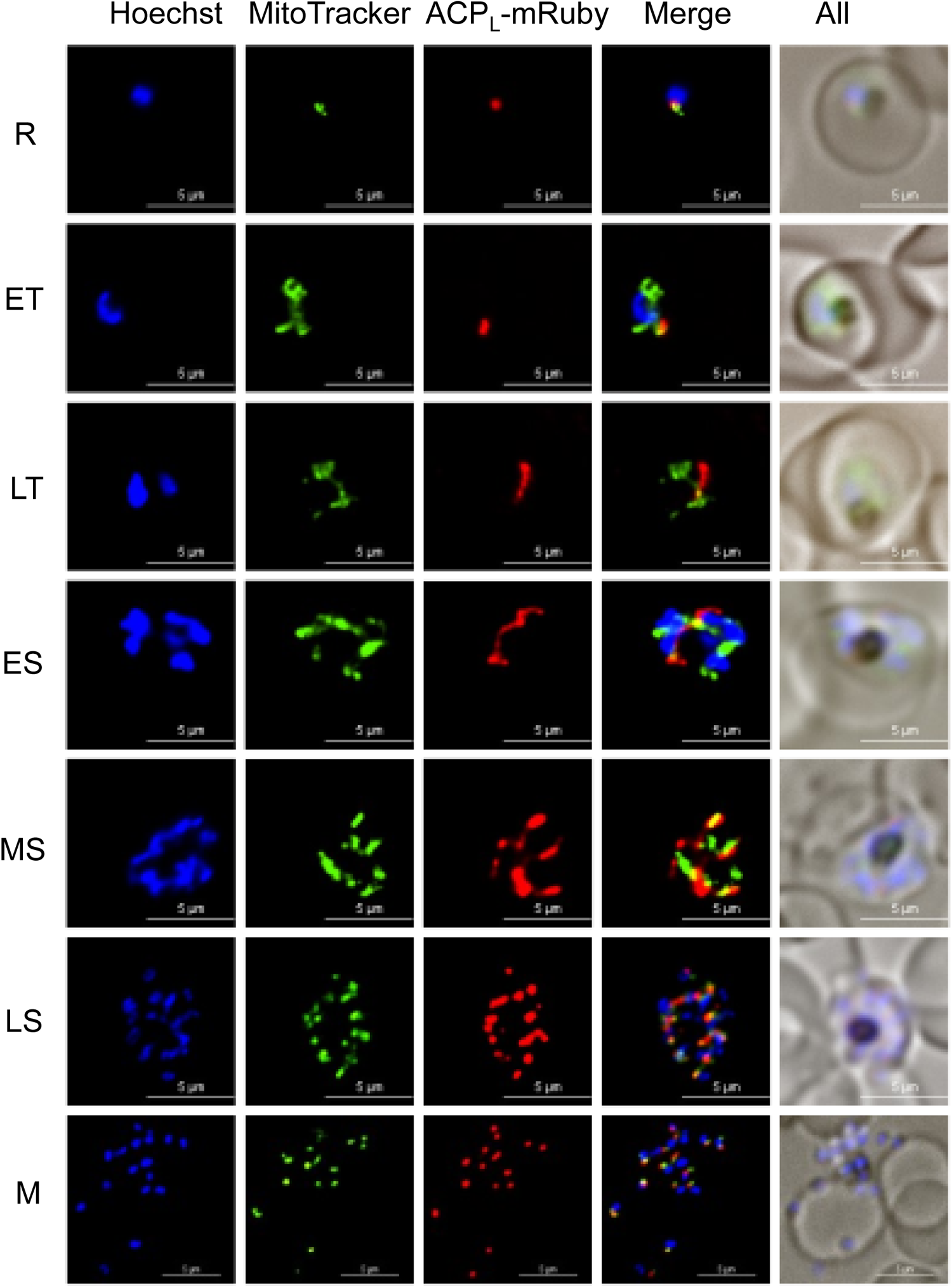
Morphologies of the apicoplast labeled with ACP_L_-mRuby in the PfDyn2 sufficient parasites. In the PfDyn2-3HA^apt^ line, the apicoplast was genetically labeled with ACP_L_-mRuby. The first 55 aa of ACP (acyl carrier protein) was infused with mRuby at its 5’ end. Mitochondria were labeled with MitoTracker Green FM (M46750, ThermoFisher). R, ring. ET, early trophozoite, LT, late trophozoite. ES, early schizont. LS, late schizont. M, merozoite. Images were taken from a Nickon Ti microscope in live conditions and processed by Nikon NIS elements software.

**Figure S6.**
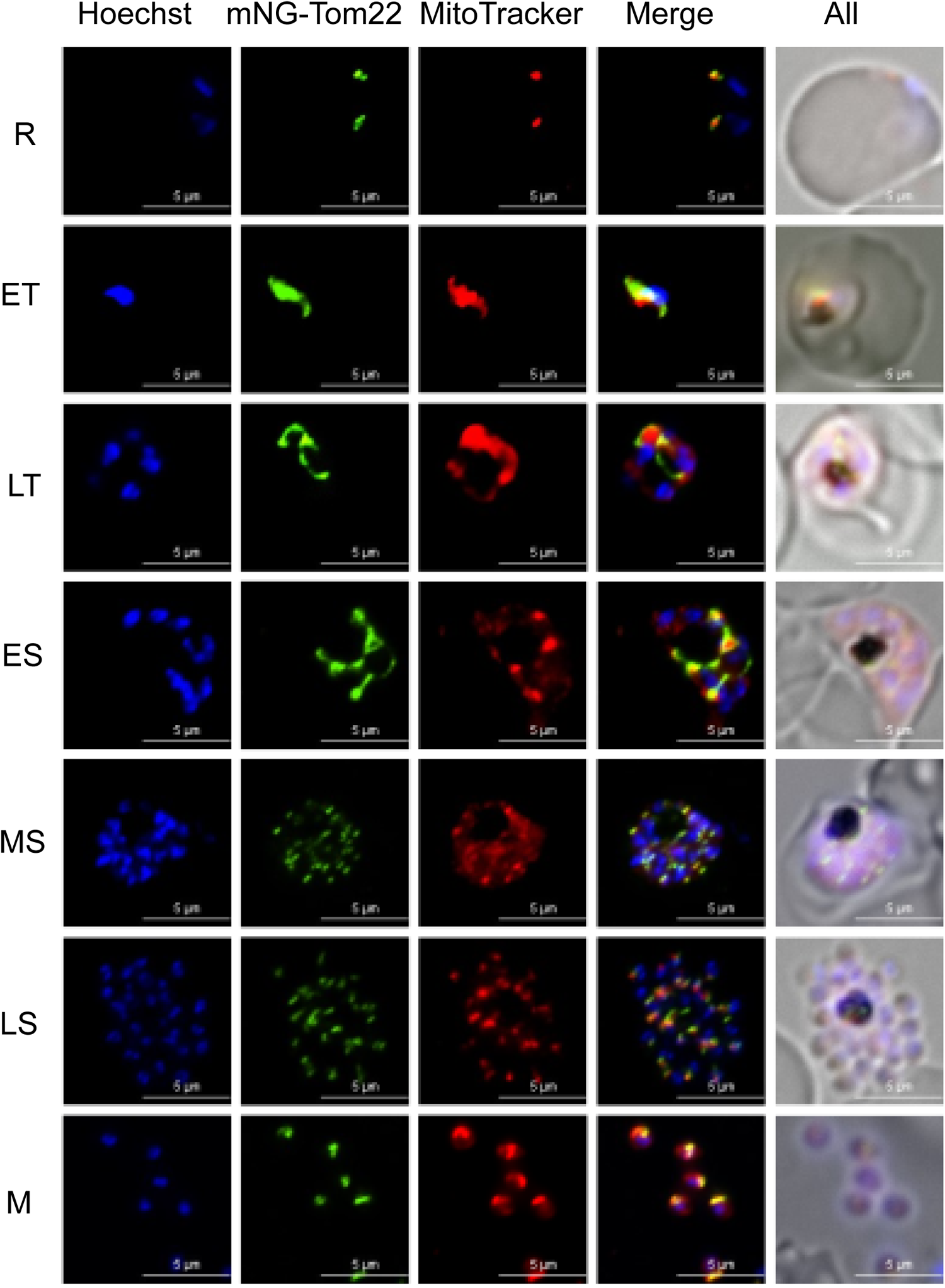
Morphologies of the mitochondria labeled with mNeonGreen in the PfDyn2 sufficient parasites. In the PfDyn2-3HA^apt^ line, the mitochondria were genetically labeled with mNeonGreen-Tom22. Mitochondria were also labeled with MitoTracker Red CMXRos. R, ring. ET, early trophozoite, LT, late trophozoite. ES, early schizont. LS, late schizont. M, merozoite. Images were taken from a Nickon Ti microscope in live conditions and processed by Nikon NIS elements software.

## Supplemental Video Legends

### Video 1

Description: 3-dimensional rendering of time-lapse confocal (AiryScan Multiplex 4Y) imaging of PfDyn2-3HA^apt^-ACP_L_-mRuby (PfDyn2-sufficient condition). ACP_L_-mRuby is shown in green alone (right panel) and merged with transmitted light / brightfield illumination of cellular morphology (left panel). Z-stacks were collected every 20 minutes for 13 hours; frame rate is 1 second per time point. Scale bar = 1 μm. Corresponds with Figure 2a (top rows).

### Video 2

Description: 3-dimensional rendering of time-lapse confocal (AiryScan Multiplex 4Y) imaging of PfDyn2-3HA^apt^-ACP_L_-mRuby parasites (PfDyn2-deficient condition). ACP_L_-mRuby is shown in green alone (right panel) and merged with transmitted light / brightfield illumination of cellular morphology (left panel). Z-stacks were collected every 20 minutes for 13 hours; frame rate is 1 second per time point. Scale bar = 1 μm. Corresponds with Figure 2a (bottom rows).

### Video 3

Description: 3-dimensional rendering of time-lapse confocal (AiryScan Multiplex 4Y) imaging of PfDyn2-3HA^apt^-StrepII-mNeonGreen-Tom22 parasites (PfDyn2-sufficient condition). mNeonGreen-Tom22 is shown in magenta alone (right panel) and merged with transmitted light / brightfield illumination of cellular morphology (left panel). Z-stacks were collected every 20 minutes for 13 hours; frame rate is 1 second per time point. Scale bar = 1 μm. Corresponds with Figure 3a (top rows).

### Video 4

Description: 3-dimensional rendering of time-lapse confocal (AiryScan Multiplex 4Y) imaging of PfDyn2-3HA^apt^-StrepII-mNeonGreen-Tom22 parasites (PfDyn2-deficient condition). mNeonGreen-Tom22 is shown in magenta alone (right panel) and merged with transmitted light / brightfield illumination of cellular morphology (left panel). Z-stacks were collected every 20 minutes for 13 hours; frame rate is 1 second per time point. Scale bar = 1 μm. Corresponds with Figure 3a (bottom rows).

### Video 5

Description: 3-dimensional rendering of time-lapse confocal (AiryScan Multiplex 4Y) imaging of PfDyn2-3HA^apt^-PfCINCH-mNeonGreen parasites (PfDyn2-sufficient condition) stained with 10 nM MitoTracker Deep Red FM. PfCINCH-mNeonGreen (green) and MitoTracker (magenta) are merged in the leftmost panel, PfCINCH alone is shown in the center panel, and MitoTracker alone is shown in the rightmost panel. Z-stacks were collected every 20 minutes for 13 hours; frame rate is 1 second per time point. Scale bar = 1 μm. Corresponds with Figure 4b (top rows).

### Video 6

Description: 3-dimensional rendering of time-lapse confocal (AiryScan Multiplex 4Y) imaging of PfDyn2-3HA^apt^-PfCINCH-mNeonGreen parasites (PfDyn2-deficient condition) stained with 10 nM MitoTracker Deep Red FM. PfCINCH-mNeonGreen (green) and MitoTracker (magenta) are merged in the leftmost panel, PfCINCH alone is shown in the center panel, and MitoTracker alone is shown in the rightmost panel. Z-stacks were collected every 20 minutes for 13 hours; frame rate is 1 second per time point. Scale bar = 1 μm. Corresponds with Figure 4b (bottom rows).

## References

1. WHO (2023). World Malaria Report.

2. van Dooren, G.G., Stimmler, L.M., and McFadden, G.I. (2006). Metabolic maps and functions of the Plasmodium mitochondrion. FEMS Microbiol Rev 30, 596–630. 10.1111/j.1574-6976.2006.00027.x.

3. Mather, M.W., Henry, K.W., and Vaidya, A.B. (2007). Mitochondrial drug targets in apicomplexan parasites. Curr Drug Targets 8, 49–60. 10.2174/138945007779315632.

4. Maclean, A.E., Hayward, J.A., Huet, D., van Dooren, G.G., and Sheiner, L. (2022). The mystery of massive mitochondrial complexes: the apicomplexan respiratory chain. Trends Parasitol 38, 1041–1052. 10.1016/j.pt.2022.09.008.

5. Hudson, A.T. (1993). Atovaquone - a novel broad-spectrum anti-infective drug. Parasitol Today 9, 66–68. 10.1016/0169-4758(93)90040-m.

6. Srivastava, I.K., Rottenberg, H., and Vaidya, A.B. (1997). Atovaquone, a broad spectrum antiparasitic drug, collapses mitochondrial membrane potential in a malarial parasite. J Biol Chem 272, 3961–3966. 10.1074/jbc.272.7.3961.

7. Looareesuwan, S., Chulay, J.D., Canfield, C.J., and Hutchinson, D.B. (1999). Malarone (atovaquone and proguanil hydrochloride): a review of its clinical development for treatment of malaria. Malarone Clinical Trials Study Group. Am J Trop Med Hyg 60, 533–541. 10.4269/ajtmh.1999.60.533.

8. McCarthy, J.S., Lotharius, J., Ruckle, T., Chalon, S., Phillips, M.A., Elliott, S., Sekuloski, S., Griffin, P., Ng, C.L., Fidock, D.A., et al. (2017). Safety, tolerability, pharmacokinetics, and activity of the novel long-acting antimalarial DSM265: a two-part first-in-human phase 1a/1b randomised study. Lancet Infect Dis 17, 626–635. 10.1016/S1473-3099(17)30171-8.

9. Sulyok, M., Ruckle, T., Roth, A., Murbeth, R.E., Chalon, S., Kerr, N., Samec, S.S., Gobeau, N., Calle, C.L., Ibanez, J., et al. (2017). DSM265 for Plasmodium falciparum chemoprophylaxis: a randomised, double blinded, phase 1 trial with controlled human malaria infection. Lancet Infect Dis 17, 636–644. 10.1016/S1473-3099(17)30139-1.

10. Llanos-Cuentas, A., Casapia, M., Chuquiyauri, R., Hinojosa, J.C., Kerr, N., Rosario, M., Toovey, S., Arch, R.H., Phillips, M.A., Rozenberg, F.D., et al. (2018). Antimalarial activity of single-dose DSM265, a novel plasmodium dihydroorotate dehydrogenase inhibitor, in patients with uncomplicated Plasmodium falciparum or Plasmodium vivax malaria infection: a proof-of-concept, open-label, phase 2a study. Lancet Infect Dis 18, 874–883. 10.1016/S1473-3099(18)30309-8.

11. Frueh, L., Li, Y., Mather, M.W., Li, Q., Pou, S., Nilsen, A., Winter, R.W., Forquer, I.P., Pershing, A.M., Xie, L.H., et al. (2017). Alkoxycarbonate Ester Prodrugs of Preclinical Drug Candidate ELQ-300 for Prophylaxis and Treatment of Malaria. ACS Infect Dis 3, 728–735. 10.1021/acsinfecdis.7b00062.

12. Smilkstein, M.J., Pou, S., Krollenbrock, A., Bleyle, L.A., Dodean, R.A., Frueh, L., Hinrichs, D.J., Li, Y., Martinson, T., Munar, M.Y., et al. (2019). ELQ-331 as a prototype for extremely durable chemoprotection against malaria. Malar J 18, 291. 10.1186/s12936-019-2921-9.

13. McFadden, G.I., Reith, M.E., Munholland, J., and Lang-Unnasch, N. (1996). Plastid in human parasites. Nature 381, 482. 10.1038/381482a0.

14. Kohler, S., Delwiche, C.F., Denny, P.W., Tilney, L.G., Webster, P., Wilson, R.J., Palmer, J.D., and Roos, D.S. (1997). A plastid of probable green algal origin in Apicomplexan parasites. Science 275, 1485–1489. 10.1126/science.275.5305.1485.

15. Janouskovec, J., Horak, A., Obornik, M., Lukes, J., and Keeling, P.J. (2010). A common red algal origin of the apicomplexan, dinoflagellate, and heterokont plastids. Proc Natl Acad Sci U S A 107, 10949–10954. 10.1073/pnas.1003335107.

16. Lemgruber, L., Kudryashev, M., Dekiwadia, C., Riglar, D.T., Baum, J., Stahlberg, H., Ralph, S.A., and Frischknecht, F. (2013). Cryo-electron tomography reveals four-membrane architecture of the Plasmodium apicoplast. Malar J 12, 25. 10.1186/1475-2875-12-25.

17. Coatney, G.R., and Greenberg, J. (1952). The use of antibiotics in the treatment of malaria. Ann N Y Acad Sci 55, 1075–1081. 10.1111/j.1749-6632.1952.tb22668.x.

18. Ralph, S.A., D’Ombrain, M.C., and McFadden, G.I. (2001). The apicoplast as an antimalarial drug target. Drug Resist Updat 4, 145–151. 10.1054/drup.2001.0205.

19. MacRae, J.I., Marechal, E., Biot, C., and Botte, C.Y. (2012). The apicoplast: a key target to cure malaria. Curr Pharm Des 18, 3490–3504.

20. McFadden, G.I., and Yeh, E. (2017). The apicoplast: now you see it, now you don’t. Int J Parasitol 47, 137–144. 10.1016/j.ijpara.2016.08.005.

21. Biddau, M., and Sheiner, L. (2019). Targeting the apicoplast in malaria. Biochem Soc Trans 47, 973–983. 10.1042/BST20170563.

22. Elahi, R., and Prigge, S.T. (2023). New insights into apicoplast metabolism in blood-stage malaria parasites. Curr Opin Microbiol 71, 102255. 10.1016/j.mib.2022.102255.

23. Painter, H.J., Morrisey, J.M., Mather, M.W., and Vaidya, A.B. (2007). Specific role of mitochondrial electron transport in blood-stage Plasmodium falciparum. Nature 446, 88–91. 10.1038/nature05572.

24. Swift, R.P., Elahi, R., Rajaram, K., Liu, H.B., and Prigge, S.T. (2023). The Plasmodium falciparum apicoplast cysteine desulfurase provides sulfur for both iron-sulfur cluster assembly and tRNA modification. Elife 12. 10.7554/eLife.84491.

25. Ke, H., Lewis, I.A., Morrisey, J.M., McLean, K.J., Ganesan, S.M., Painter, H.J., Mather, M.W., Jacobs-Lorena, M., Llinas, M., and Vaidya, A.B. (2015). Genetic investigation of tricarboxylic acid metabolism during the Plasmodium falciparum life cycle. Cell Rep 11, 164–174. 10.1016/j.celrep.2015.03.011.

26. Ke, H., Sigala, P.A., Miura, K., Morrisey, J.M., Mather, M.W., Crowley, J.R., Henderson, J.P., Goldberg, D.E., Long, C.A., and Vaidya, A.B. (2014). The heme biosynthesis pathway is essential for Plasmodium falciparum development in mosquito stage but not in blood stages. J Biol Chem 289, 34827–34837. 10.1074/jbc.M114.615831.

27. Shunmugam, S., Arnold, C.S., Dass, S., Katris, N.J., and Botte, C.Y. (2022). The flexibility of Apicomplexa parasites in lipid metabolism. PLoS Pathog 18, e1010313. 10.1371/journal.ppat.1010313.

28. Yeh, E., and DeRisi, J.L. (2011). Chemical rescue of malaria parasites lacking an apicoplast defines organelle function in blood-stage Plasmodium falciparum. PLoS Biol 9, e1001138. 10.1371/journal.pbio.1001138.

29. Swift, R.P., Rajaram, K., Liu, H.B., and Prigge, S.T. (2021). Dephospho-CoA kinase, a nuclear-encoded apicoplast protein, remains active and essential after Plasmodium falciparum apicoplast disruption. EMBO J 40, e107247. 10.15252/embj.2020107247.

30. van Dooren, G.G., Marti, M., Tonkin, C.J., Stimmler, L.M., Cowman, A.F., and McFadden, G.I. (2005). Development of the endoplasmic reticulum, mitochondrion and apicoplast during the asexual life cycle of Plasmodium falciparum. Mol Microbiol 57, 405–419. 10.1111/j.1365-2958.2005.04699.x.

31. Verhoef, J.M.J., Meissner, M., and Kooij, T.W.A. (2021). Organelle Dynamics in Apicomplexan Parasites. mBio 12, e0140921. 10.1128/mBio.01409-21.

32. Elaagip, A., Absalon, S., and Florentin, A. (2022). Apicoplast Dynamics During Plasmodium Cell Cycle. Front Cell Infect Microbiol 12, 864819. 10.3389/fcimb.2022.864819.

33. Morano, A.A., and Dvorin, J.D. (2021). The Ringleaders: Understanding the Apicomplexan Basal Complex Through Comparison to Established Contractile Ring Systems. Front Cell Infect Microbiol 11, 656976. 10.3389/fcimb.2021.656976.

34. Voss, Y., Klaus, S., Guizetti, J., and Ganter, M. (2023). Plasmodium schizogony, a chronology of the parasite’s cell cycle in the blood stage. PLoS Pathog 19, e1011157. 10.1371/journal.ppat.1011157.

35. Rudlaff, R.M., Kraemer, S., Marshman, J., and Dvorin, J.D. (2020). Three-dimensional ultrastructure of Plasmodium falciparum throughout cytokinesis. PLoS Pathog 16, e1008587. 10.1371/journal.ppat.1008587.

36. Verhoef, J.M.J., Boshoven, C., Evers, F., Akkerman, L.J., Gijsbrechts, B.C.A., van de Vegte-Bolmer, M., van Gemert, G.J., Vaidya, A.B., and Kooij, T.W.A. (2024). Detailing organelle division and segregation in Plasmodium falciparum. bioRxiv. 10.1101/2024.01.30.577899.

37. Ferguson, S.M., and De Camilli, P. (2012). Dynamin, a membrane-remodelling GTPase. Nat Rev Mol Cell Biol 13, 75–88. 10.1038/nrm3266.

38. Ford, M.G., Jenni, S., and Nunnari, J. (2011). The crystal structure of dynamin. Nature 477, 561–566. 10.1038/nature10441.

39. van Dooren, G.G., Reiff, S.B., Tomova, C., Meissner, M., Humbel, B.M., and Striepen, B. (2009). A novel dynamin-related protein has been recruited for apicoplast fission in Toxoplasma gondii. Curr Biol 19, 267–276. 10.1016/j.cub.2008.12.048.

40. Breinich, M.S., Ferguson, D.J., Foth, B.J., van Dooren, G.G., Lebrun, M., Quon, D.V., Striepen, B., Bradley, P.J., Frischknecht, F., Carruthers, V.B., and Meissner, M. (2009). A dynamin is required for the biogenesis of secretory organelles in Toxoplasma gondii. Curr Biol 19, 277–286. 10.1016/j.cub.2009.01.039.

41. Heredero-Bermejo, I., Varberg, J.M., Charvat, R., Jacobs, K., Garbuz, T., Sullivan, W.J., Jr., and Arrizabalaga, G. (2019). TgDrpC, an atypical dynamin-related protein in Toxoplasma gondii, is associated with vesicular transport factors and parasite division. Mol Microbiol 111, 46–64. 10.1111/mmi.14138.

42. Melatti, C., Pieperhoff, M., Lemgruber, L., Pohl, E., Sheiner, L., and Meissner, M. (2019). A unique dynamin-related protein is essential for mitochondrial fission in Toxoplasma gondii. PLoS Pathog 15, e1007512. 10.1371/journal.ppat.1007512.

43. Zhou, H.C., Gao, Y.H., Zhong, X., and Wang, H. (2009). Dynamin like protein 1 participated in the hemoglobin uptake pathway of Plasmodium falciparum. Chin Med J (Engl) 122, 1686–1691.

44. Li, H., Han, Z., Lu, Y., Lin, Y., Zhang, L., Wu, Y., and Wang, H. (2004). Isolation and functional characterization of a dynamin-like gene from Plasmodium falciparum. Biochem Biophys Res Commun 320, 664–671. 10.1016/j.bbrc.2004.06.010.

45. Charneau, S., Bastos, I.M., Mouray, E., Ribeiro, B.M., Santana, J.M., Grellier, P., and Florent, I. (2007). Characterization of PfDYN2, a dynamin-like protein of Plasmodium falciparum expressed in schizonts. Microbes Infect 9, 797–805. 10.1016/j.micinf.2007.02.020.

46. Ganesan, S.M., Falla, A., Goldfless, S.J., Nasamu, A.S., and Niles, J.C. (2016). Synthetic RNA-protein modules integrated with native translation mechanisms to control gene expression in malaria parasites. Nat Commun 7, 10727. 10.1038/ncomms10727.

47. Rajaram, K., Liu, H.B., and Prigge, S.T. (2020). Redesigned TetR-Aptamer System To Control Gene Expression in Plasmodium falciparum. mSphere 5. 10.1128/mSphere.00457-20.

48. Gambarotto, D., Zwettler, F.U., Le Guennec, M., Schmidt-Cernohorska, M., Fortun, D., Borgers, S., Heine, J., Schloetel, J.G., Reuss, M., Unser, M., et al. (2019). Imaging cellular ultrastructures using expansion microscopy (U-ExM). Nat Methods 16, 71–74. 10.1038/s41592-018-0238-1.

49. Liffner, B., and Absalon, S. (2021). Expansion Microscopy Reveals Plasmodium falciparum Blood-Stage Parasites Undergo Anaphase with A Chromatin Bridge in the Absence of Mini-Chromosome Maintenance Complex Binding Protein. Microorganisms 9. 10.3390/microorganisms9112306.

50. Gisselberg, J.E., Dellibovi-Ragheb, T.A., Matthews, K.A., Bosch, G., and Prigge, S.T. (2013). The suf iron-sulfur cluster synthesis pathway is required for apicoplast maintenance in malaria parasites. PLoS Pathog 9, e1003655. 10.1371/journal.ppat.1003655.

51. Pasaje, C.F., Cheung, V., Kennedy, K., Lim, E.E., Baell, J.B., Griffin, M.D., and Ralph, S.A. (2016). Selective inhibition of apicoplast tryptophanyl-tRNA synthetase causes delayed death in Plasmodium falciparum. Sci Rep 6, 27531. 10.1038/srep27531.

52. Florentin, A., Cobb, D.W., Fishburn, J.D., Cipriano, M.J., Kim, P.S., Fierro, M.A., Striepen, B., and Muralidharan, V. (2017). PfClpC Is an Essential Clp Chaperone Required for Plastid Integrity and Clp Protease Stability in Plasmodium falciparum. Cell Rep 21, 1746–1756. 10.1016/j.celrep.2017.10.081.

53. Uddin, T., McFadden, G.I., and Goodman, C.D. (2018). Validation of Putative Apicoplast-Targeting Drugs Using a Chemical Supplementation Assay in Cultured Human Malaria Parasites. Antimicrob Agents Chemother 62. 10.1128/AAC.01161-17.

54. Walczak, M., Ganesan, S.M., Niles, J.C., and Yeh, E. (2018). ATG8 Is Essential Specifically for an Autophagy-Independent Function in Apicoplast Biogenesis in Blood-Stage Malaria Parasites. mBio 9. 10.1128/mBio.02021-17.

55. Sayers, C.P., Mollard, V., Buchanan, H.D., McFadden, G.I., and Goodman, C.D. (2018). A genetic screen in rodent malaria parasites identifies five new apicoplast putative membrane transporters, one of which is essential in human malaria parasites. Cell Microbiol 20. 10.1111/cmi.12789.

56. Tang, Y., Meister, T.R., Walczak, M., Pulkoski-Gross, M.J., Hari, S.B., Sauer, R.T., Amberg-Johnson, K., and Yeh, E. (2019). A mutagenesis screen for essential plastid biogenesis genes in human malaria parasites. PLoS Biol 17, e3000136. 10.1371/journal.pbio.3000136.

57. Meister, T.R., Tang, Y., Pulkoski-Gross, M.J., and Yeh, E. (2020). CaaX-Like Protease of Cyanobacterial Origin Is Required for Complex Plastid Biogenesis in Malaria Parasites. mBio 11. 10.1128/mBio.01492-20.

58. Swift, R.P., Rajaram, K., Keutcha, C., Liu, H.B., Kwan, B., Dziedzic, A., Jedlicka, A.E., and Prigge, S.T. (2020). The NTP generating activity of pyruvate kinase II is critical for apicoplast maintenance in Plasmodium falciparum. Elife 9. 10.7554/eLife.50807.

59. Tan, S., Mudeppa, D.G., Kokkonda, S., White, J., 3rd, Patrapuvich, R., and Rathod, P.K. (2021). Properties of Plasmodium falciparum with a Deleted Apicoplast DNA Gyrase. Antimicrob Agents Chemother 65, e0058621. 10.1128/AAC.00586-21.

60. Okada, M., Rajaram, K., Swift, R.P., Mixon, A., Maschek, J.A., Prigge, S.T., and Sigala, P.A. (2022). Critical role for isoprenoids in apicoplast biogenesis by malaria parasites. Elife 11. 10.7554/eLife.73208.

61. Nkrumah, L.J., Muhle, R.A., Moura, P.A., Ghosh, P., Hatfull, G.F., Jacobs, W.R., Jr., and Fidock, D.A. (2006). Efficient site-specific integration in Plasmodium falciparum chromosomes mediated by mycobacteriophage Bxb1 integrase. Nat Methods 3, 615–621. 10.1038/nmeth904.

62. Gallagher, J.R., Matthews, K.A., and Prigge, S.T. (2011). Plasmodium falciparum apicoplast transit peptides are unstructured in vitro and during apicoplast import. Traffic 12, 1124–1138. 10.1111/j.1600-0854.2011.01232.x.

63. Hale, V.L., Watermeyer, J.M., Hackett, F., Vizcay-Barrena, G., van Ooij, C., Thomas, J.A., Spink, M.C., Harkiolaki, M., Duke, E., Fleck, R.A., et al. (2017). Parasitophorous vacuole poration precedes its rupture and rapid host erythrocyte cytoskeleton collapse in Plasmodium falciparum egress. Proc Natl Acad Sci U S A 114, 3439–3444. 10.1073/pnas.1619441114.

64. Holder, A.A. (2009). The carboxy-terminus of merozoite surface protein 1: structure, specific antibodies and immunity to malaria. Parasitology 136, 1445–1456. 10.1017/S0031182009990515.

65. Gubbels, M.J., Ferguson, D.J.P., Saha, S., Romano, J.D., Chavan, S., Primo, V.A., Michaud, C., Coppens, I., and Engelberg, K. (2022). Toxoplasma gondii’s Basal Complex: The Other Apicomplexan Business End Is Multifunctional. Front Cell Infect Microbiol 12, 882166. 10.3389/fcimb.2022.882166.

66. Cepeda Diaz, A.K., Rudlaff, R.M., Farringer, M., and Dvorin, J.D. (2023). Essential function of alveolin PfIMC1g in the Plasmodium falciparum asexual blood stage. mBio 14, e0150723. 10.1128/mbio.01507-23.

67. Rudlaff, R.M., Kraemer, S., Streva, V.A., and Dvorin, J.D. (2019). An essential contractile ring protein controls cell division in Plasmodium falciparum. Nat Commun 10, 2181. 10.1038/s41467-019-10214-z.

68. Milani, K.J., Schneider, T.G., and Taraschi, T.F. (2015). Defining the morphology and mechanism of the hemoglobin transport pathway in Plasmodium falciparum-infected erythrocytes. Eukaryot Cell 14, 415–426. 10.1128/EC.00267-14.

69. Morano, A.A., Rudlaff, R.M., and Dvorin, J.D. (2023). A PPP-type pseudophosphatase is required for the maintenance of basal complex integrity in Plasmodium falciparum. Nat Commun 14, 3916. 10.1038/s41467-023-39435-z.

70. McDonald, B., and Martin-Serrano, J. (2009). No strings attached: the ESCRT machinery in viral budding and cytokinesis. J Cell Sci 122, 2167–2177. 10.1242/jcs.028308.

71. Caballe, A., and Martin-Serrano, J. (2011). ESCRT machinery and cytokinesis: the road to daughter cell separation. Traffic 12, 1318–1326. 10.1111/j.1600-0854.2011.01244.x.

72. Leung, K.F., Dacks, J.B., and Field, M.C. (2008). Evolution of the multivesicular body ESCRT machinery; retention across the eukaryotic lineage. Traffic 9, 1698–1716. 10.1111/j.1600-0854.2008.00797.x.

73. Maruthi, M., Ling, L., Zhou, J., and Ke, H. (2020). Dispensable Role of Mitochondrial Fission Protein 1 (Fis1) in the Erythrocytic Development of Plasmodium falciparum. mSphere 5. 10.1128/mSphere.00579-20.

74. Jacobs, K., Charvat, R., and Arrizabalaga, G. (2020). Identification of Fis1 Interactors in Toxoplasma gondii Reveals a Novel Protein Required for Peripheral Distribution of the Mitochondrion. mBio 11. 10.1128/mBio.02732-19.

75. Das, S., Lemgruber, L., Tay, C.L., Baum, J., and Meissner, M. (2017). Multiple essential functions of Plasmodium falciparum actin-1 during malaria blood-stage development. BMC Biol 15, 70. 10.1186/s12915-017-0406-2.

76. M’Saad, O., and Bewersdorf, J. (2020). Light microscopy of proteins in their ultrastructural context. Nat Commun 11, 3850. 10.1038/s41467-020-17523-8.

77. Lorestani, A., Sheiner, L., Yang, K., Robertson, S.D., Sahoo, N., Brooks, C.F., Ferguson, D.J., Striepen, B., and Gubbels, M.J. (2010). A Toxoplasma MORN1 null mutant undergoes repeated divisions but is defective in basal assembly, apicoplast division and cytokinesis. PLoS One 5, e12302. 10.1371/journal.pone.0012302.

78. Heaslip, A.T., Dzierszinski, F., Stein, B., and Hu, K. (2010). TgMORN1 is a key organizer for the basal complex of Toxoplasma gondii. PLoS Pathog 6, e1000754. 10.1371/journal.ppat.1000754.

79. Oliveira Souza, R.O., Jacobs, K.N., Back, P.S., Bradley, P.J., and Arrizabalaga, G. (2022). IMC10 and LMF1 mediate mitochondrial morphology through mitochondrion-pellicle contact sites in Toxoplasma gondii. J Cell Sci 135. 10.1242/jcs.260083.

80. Striepen, B., Crawford, M.J., Shaw, M.K., Tilney, L.G., Seeber, F., and Roos, D.S. (2000). The plastid of Toxoplasma gondii is divided by association with the centrosomes. J Cell Biol 151, 1423–1434. 10.1083/jcb.151.7.1423.

81. Ovciarikova, J., Lemgruber, L., Stilger, K.L., Sullivan, W.J., and Sheiner, L. (2017). Mitochondrial behaviour throughout the lytic cycle of Toxoplasma gondii. Sci Rep 7, 42746. 10.1038/srep42746.

82. Liffner, B., Cepeda Diaz, A.K., Blauwkamp, J., Anaguano, D., Frolich, S., Muralidharan, V., Wilson, D.W., Dvorin, J., and Absalon, S. (2023). Atlas of Plasmodium falciparum intraerythrocytic development using expansion microscopy. bioRxiv. 10.1101/2023.03.22.533773.

83. Ke, H., Dass, S., Morrisey, J.M., Mather, M.W., and Vaidya, A.B. (2018). The mitochondrial ribosomal protein L13 is critical for the structural and functional integrity of the mitochondrion in Plasmodium falciparum. J Biol Chem 293, 8128–8137. 10.1074/jbc.RA118.002552.

84. Ling, L., Mulaka, M., Munro, J., Dass, S., Mather, M.W., Riscoe, M.K., Llinas, M., Zhou, J., and Ke, H. (2020). Genetic ablation of the mitoribosome in the malaria parasite Plasmodium falciparum sensitizes it to antimalarials that target mitochondrial functions. J Biol Chem 295, 7235–7248. 10.1074/jbc.RA120.012646.

85. Solebo, O., Ling, L., Nwankwo, I., Zhou, J., Fu, T.M., and Ke, H. (2023). Plasmodium falciparum utilizes pyrophosphate to fuel an essential proton pump in the ring stage and the transition to trophozoite stage. PLoS Pathog 19, e1011818. 10.1371/journal.ppat.1011818.

## References

1. Ganesan, S.M., Falla, A., Goldfless, S.J., Nasamu, A.S., and Niles, J.C. (2016). Synthetic RNA-protein modules integrated with native translation mechanisms to control gene expression in malaria parasites. Nat Commun 7, 10727. 10.1038/ncomms10727.

2. Rajaram, K., Liu, H.B., and Prigge, S.T. (2020). Redesigned TetR-Aptamer System To Control Gene Expression in Plasmodium falciparum. mSphere 5. 10.1128/mSphere.00457-20.

3. Solebo, O., Ling, L., Nwankwo, I., Zhou, J., Fu, T.M., and Ke, H. (2023). Plasmodium falciparum utilizes pyrophosphate to fuel an essential proton pump in the ring stage and the transition to trophozoite stage. PLoS Pathog 19, e1011818. 10.1371/journal.ppat.1011818.

4. Ling, L., Mulaka, M., Munro, J., Dass, S., Mather, M.W., Riscoe, M.K., Llinas, M., Zhou, J., and Ke, H. (2020). Genetic ablation of the mitoribosome in the malaria parasite Plasmodium falciparum sensitizes it to antimalarials that target mitochondrial functions. J Biol Chem 295, 7235–7248. 10.1074/jbc.RA120.012646.

5. Lamb, I.M., Rios, K.T., Shukla, A., Ahiya, A.I., Morrisey, J., Mell, J.C., Lindner, S.E., Mather, M.W., and Vaidya, A.B. (2022). Mitochondrially targeted proximity biotinylation and proteomic analysis in Plasmodium falciparum. PLoS One 17, e0273357. 10.1371/journal.pone.0273357.

6. Ke, H., Morrisey, J.M., Ganesan, S.M., Painter, H.J., Mather, M.W., and Vaidya, A.B. (2011). Variation among Plasmodium falciparum strains in their reliance on mitochondrial electron transport chain function. Eukaryot Cell 10, 1053–1061. 10.1128/EC.05049-11.

7. van Dooren, G.G., Yeoh, L.M., Striepen, B., and McFadden, G.I. (2016). The Import of Proteins into the Mitochondrion of Toxoplasma gondii. J Biol Chem 291, 19335–19350. 10.1074/jbc.M116.725069.

8. Ke, H., Dass, S., Morrisey, J.M., Mather, M.W., and Vaidya, A.B. (2018). The mitochondrial ribosomal protein L13 is critical for the structural and functional integrity of the mitochondrion in Plasmodium falciparum. J Biol Chem 293, 8128–8137. 10.1074/jbc.RA118.002552.

9. Liffner, B., and Absalon, S. (2021). Expansion Microscopy Reveals Plasmodium falciparum Blood-Stage Parasites Undergo Anaphase with A Chromatin Bridge in the Absence of Mini-Chromosome Maintenance Complex Binding Protein. Microorganisms 9. 10.3390/microorganisms9112306.

